# Pith cell responses to low red: far-red light in dicot stems

**DOI:** 10.1101/2024.11.12.623171

**Authors:** Linge Li, Yorrit van de Kaa, Lotte van der Krabben, Ronald Pierik, Kaisa Kajala

**Author notes:** **Email address for each author:**.

## Abstract

In dense canopies light becomes a limiting factor for plant growth. Many plants respond to neighbor cues by growing taller to improve light capture, a phenomenon known as the shade avoidance syndrome (SAS). The major neighbor detection is via enrichment of far-red (FR) light that leads to a low red : far-red light ratio (R:FR), suppressing phytochrome activity. In tomato, low R:FR induces elongation of the internodes, but study into the role of different cell types in this response has remained limited.

We characterized changes in cellular anatomy of the tomato internode in response to low R:FR, and its accompanying changes in gene expression. We observed changes to the pith traits, including increases in pith layer number, pith cell diameter and longitudinal cell length. We profiled the transcriptome in the entire internodes and in the hand-dissected pith in the central cylinder of the internode in response to low R:FR treatment and identified transcription factors (TFs) of interest that were upregulated in the central cylinder, mostly GATA, TCP, and bZIPs. We then characterized FR-responses in eight dicotyledonous species. Significant pith elongation was observed in species that exhibited a strong internode elongation response. The FR-responsive expression of homologs of target GATA, TCP and bZIP TFs in the central cylinder was conserved within the Solanaceae family.

Overall, we discovered central cylinder gene expression patterns in SAS that are distinct from those of the entire internode, suggesting that some responses are unique and likely specific to vascular cell types such as pith. These patterns were conserved with close relatives of tomato but not in other dicot families we sampled, indicating that different molecular mechanisms drive FR responses in different dicots.

## Introduction

The challenge of meeting the growing demands for food, fuel, and fiber in the face of increasing population and limited land area has led to the intensive cultivation of plants in dense vegetation. This approach, while maximizing the use of available land, introduces a significant problem: competition for limited resources, particularly light. Dense vegetation often triggers a phenomenon known as Shade Avoidance Syndrome (SAS), where plants compete for light, leading to various morphological adaptations most commonly making the plant taller and more erect (Smith and Whitelam, 1997). SAS is usually triggered by one of two common light cues: either by the enrichment of far-red (FR) light in comparison to red (R) light as only FR is reflected from and passed through the neighbouring or shading leaves, or by depletion of blue light, indicating overall reduction of light intensity. Specifically, FR enrichment is detected through the ratio of red (R) to far-red (FR) light by phytochromes. The shade-induced changes, including stem elongation and altered leaf architecture, have been observed across various plant species, such as Arabidopsis, *Brassica rapa*, and tomato (Osborne, 1991; Pierik and De Wit, 2014; Kohnen *et al*., 2016; Ballaré and Pierik, 2017).

However, SAS is not the sole strategy for plant species to deal with low light. Exploring the strategies of plant species in different ecosystems reveals approaches contrasting to competing for light via SAS. Pioneer species and shade-tolerant species demonstrate unique life strategies and can provide evolutionary perspectives on how plants respond to shade. These species exemplify the diverse ways plants thrive in various environments. Pioneer plants are equipped to establish themselves in harsh, sterile, and sun-exposed environments, rapidly growing to utilize available resources before larger competitors can establish dominance. (Miyazawa *et al*., 2014; Sottosanti, 2023). However, as the environment changes and the intermediate species grow taller, the shade they cast can deprive the pioneer species of adequate sunlight (Sottosanti, 2023). In response to shading, the composition of species transitions from pioneer plants to a mixed ecosystem. This ecosystem includes dominant plants that prefer high light and understory species that flourish in the shade provided by the forest canopy (Hagen *et al*., 2015; Avalos and Avalos, 2019). How these different light-use strategies, especially shade avoidance and tolerance, have evolved is not well characterized. SAS may have evolved alongside shade itself, potentially as early as the Late Devonian period, suggesting it could be more ancient than shade tolerance mechanisms in plants. (Mathews, 2006). In contrast, the evolution of shade tolerance is associated with attenuation of shade avoidance and reduced phenotypic plasticity in some dicot lineages such as milkweeds and *Geranium robertianum* (Gommers *et al*., 2017; Coverdale and Agrawal, 2021). Therefore, there is evidence that in these lineages, SAS predates the evolution of shade tolerance. It would be reasonable to assume that SAS mechanisms are conserved between angiosperms, but this has not been extensively tested.

A lot of SAS research has been carried out in the model species *Arabidopsis thaliana* with a focus on cellular events in the epidermis as a driver of the petiole responses (de Wit *et al*., 2012; Pierik and De Wit, 2014; van Gelderen *et al*., 2018). The Arabidopsis and its rosette growth habit however have their limitations, including not being a good model for stem elongation as a SAS trait. In this study, we set out instead to investigate how tomato (*Solanum lycopersicum*) and specifically its internodes respond to low R:FR. We paired organ-level measurements with cellular anatomy and transcriptomic data. We discovered changes in the pith cell length and layer number and accompanying transcript changes in response to low R:FR. Pith is a tissue located in the central cylinder of stems and composed of undifferentiated parenchyma cells (Zabel and Morrell, 2020), currently hypothesized to have evolved through delayed and shortened protoxylem differentiation during early euphyllophyte evolution by Middle Devonian period (Tomescu and Mcqueen, 2022). Hence, it is possible that pith evolution preceded that of SAS, so here, we tested the hypothesis that pith plays a key role in internode elongation. Thus, we queried if pith morphological and transcriptomic responses to low R:FR were conserved within dicots of increasing distance to tomato. We assessed species from the Solanaceae, Brassicaceae, and Fabaceae and found some conservation in pith responses.

From the transcriptomics data, we identified three candidate transcription factors (TF) that responded to low R:FR in tomato internodes or in their pith. These were a GATA TF (*Solyc01g0S07C0*), a TCP TF (*Solyc08g080150* or *SlTCP20*), and a bZIP TF (*Solyc07g053450*). We tested if their FR-responsive behavior was conserved in our set of species and discovered Solanaceae-specific conservation in these SAS responses. These family-specific responses to low R:FR indicate that there is divergence between upregulated TFs and molecular mechanisms involved in stem elongation during SAS of different dicots. This highlights the need for diverse model species for understanding seemingly conserved plant responses to the environment.

## Materials G methods

### Plant materials and growth conditions

We germinated seeds of *Solanum lycopersicum* (cv Moneymaker (obtained from Intratuin B.V) and M82 (obtained from Tomato Genetics Resource Center)), *Capsicum annuum*, *Solanum melongena* (obtained from Intratuin B.V), *Pisum sativum* Cameor (obtained from Dr. Judith Burstin lab), and *Glycine max* (cv green shell) (obtained from https://www.dehobbytuinder.nl/) by placing them in sealed plastic containers lined with paper towels soaked in tap water for one week. After this germination period, we transplanted seedlings of uniform size into 7 cm square pots filled with sieved Primasta® soil. Approximately one week following the transfer, we randomly allocated the plants into two distinct groups and subjected them to either white light (WL) or supplemental far-red light (WL+FR) treatments as described in Li *et al*., 2024. The seedlings were cultivated at a temperature of 20°C, maintaining a humidity level of 70%, under a 16-hour light/8-hour dark cycle. We maintained the PAR value at 200 µmol/m²/s for both WL and WL+FR. For the WL+FR, we added FR light to the same PAR background to achieve a specific R:FR ratio of 0.2.

We sowed *Arabidopsis thaliana* (Colombia-0 (Col-0)) and *Brassica nigra* seeds (Pantazopoulou *et al*., 2017) directly into the soil, treated them with a stratification period of 3 days in darkness at 4°C, and subsequently transferred them to the following conditions: LD: 16 hours of light, 8 hours of darkness, SD: 8 hours of light, 11 hours of darkness photosynthetically active radiation (PAR) of 150 μmol·m^−2^·s^−1^, an R/FR ratio of 2.0, a temperature of 20°C, and a relative humidity of 70%. Following a 7-day growth period, we transplanted the seedlings into 70 mL pots containing Primasta ® soil. At the age of approximately 2 months, we used Col-0 plants in inflorescence stem elongation experiments, commencing when the inflorescence reached approximately 0.5 cm to 2 cm in length.

All experimental procedures were initiated at ZT = 3.5 h.

### Phenotyping

We used a digital calliper to measure specific plant attributes accurately. For each species, we had 8 replicates, and the experiments were conducted twice for consistency. In the case of Arabidopsis, the measurements were carried out with 12 replicates and the experiment was repeated twice. The key phenotype measurements, specifically stem and internode length, were recorded every day at two fixed times, ZT=3.5h and ZT=9.5.

### RNA-seq

We harvested the first internode, located between the cotyledons and the first true leaf. We gathered groups of these internodes at intervals of 6, 24, 30, and 48 hours following the commencement of the treatment at ZT = 3.5 h. Ultimately, this resulted in the collection of samples encompassing two lighting conditions (WL and WL+FR), two different cultivars (M82 and Moneymaker), four timepoints, and two types of tissues (internode and its central cylinder), each with a minimum of four biological replicates. Each biological replicate contained 6-8 individual internodes or central cylinders of the internode. To preserve their integrity for mRNA isolation, the samples were promptly frozen in liquid nitrogen. RNA isolation, library construction and high-throughput sequencing were carried out exactly as we described in Li *et al*., 2024. Similarly, the analysis of the RNA-seq (mapping and differential expression analysis) was conducted as we described in Li *et al*., 2024. Raw counts are available on NCBI GEO (GSE255611), and DEGs and enriched GO terms are listed in Tables S1-S2. In our RNA-seq analysis, we compared the M82 and MM tomato cultivars to identify intrinsic differences between these varieties (Figure S1). Our results did not reveal significant variations, with the exception of minor discrepancies observed in the central cylinder data. Therefore, for other analyses, we combined these two cultivars and use it as one species to further investigate species differences.

### Histology

We stained the sections form the middle 1cm of the first internode. We immersed the internode section into 0.1% (w/v) safranin solution was two minutes. Following this, we rinsed the samples with 70% ethanol repeatedly until the ethanol appeared clear. For long-term preservation, we placed the samples back into 70% ethanol solution. We prepared sections of internode tissue with a vibratome (LEICA VT1000S) and imaged the sections with the Leica 700 Microscope with analyzeD software. We then measured specific parameters with ImageJ software as listed in Figure 1. All images were taken at a magnification of 10X. We used GraphPad Prism 9 tor the analysis and presentation of data. Furthermore, statistical analyses, including the use of the Student t-test, were performed with Microsoft Excel.

**Figure 1:**
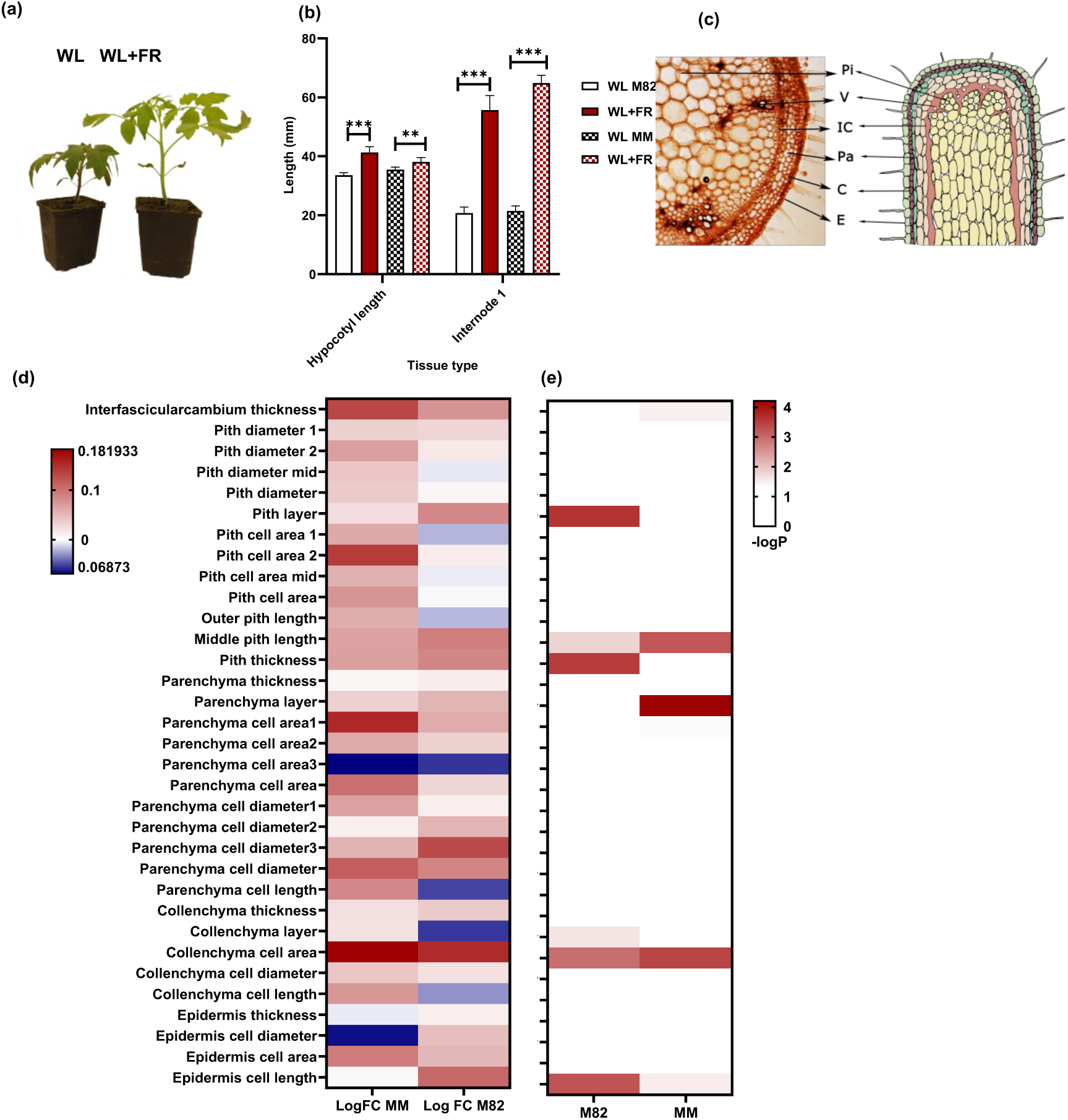
Tomato internode cellular morphology in response to low R:FR. a. Comparison of *S. lycopersicum* cultivars under different light conditions. Two 24-day-old M82 plants after 10 days of treatment of either white light (WL) or white light supplemented with far-red (WL+FR). b. Stem length analysis in M82 and Moneymaker (MM) of three-week-old plants following seven-day treatments under WL and WL+FR. Notable differences between WL and FR+WL conditions are indicated with asterisks (* p≤0.05; ** p≤0.01; *** p≤0.001). This analysis was conducted with 18 biological replicates and repeated three times. MM data are from Li *et al*., 2024. c. Cellular anatomy of M82 first internode. The cross-section details the cell types of interest including Epidermis (E), Collenchyma (C), Parenchyma (Pa), Interfascicular Cambium (IC), Vasculature (V, combined xylem and phloem) and Pith (Pi) d, e. FR-responsive changes in cellular anatomy traits in the tomato first internode. The heatmaps show (d) logFC and (e) −log(p-value) for each trait comparing WL+FR to WL. The layer number of each cell type is indicated from the outermost layer moving inward. Measurements of interfascicular cambium thickness were done in areas without vascular bundles. A logFC (logarithm of fold-change) of 0 is depicted in white to signify no change, with upregulation shown in red and downregulation in blue. A −log Pval of −log0.05 is set to white, indicating no statistical significance.

### Promoter cloning and recombination

Promoter sequences were synthesized by GenScript (Table S3). Upon synthesis, we cloned these promoters into pENTR-D/TOPO vectors (Invitrogen) through a topoisomerase I-mediated cloning process and subsequently introduced into PMR074 destination vectors (Ron *et al*., 2014). This recombination was performed using the LR Clonase II Enzyme Mix (Invitrogen), following the manufacturer’s protocol for LR recombination. Each cloning and recombination step was verified by colony PCR and subsequent sequencing to confirm the integrity and correct orientation of the inserted promoter sequences. Final binary vector for plant transformation was then transformed into *Agrobacterium tumefaciens* AGL1 cells following protocol by Liu *et al*., (2023*a*), using Rif20/Strep50 selection and a colony PCR for verification.

### Tomato transformation

We sterilized the seeds of *Solanum lycopersicum* Moneymaker by immersing them briefly in 70% ethanol followed by a 20-minute dip in a 2.5% sodium hypochlorite solution. After multiple rinses with deionized water, we placed the seeds on a hormone-free germination medium within sterile conditions and germinated them under long-day conditions at 20°C and 70% relative humidity.

The germination medium (GM) we used was composed of 0.49% Murashige and Skoog (MS) basal medium supplemented with vitamins, 2-Morpholinoethanesulfonic acid (MES), 2% sucrose, and 0.8% plant agar, pH 5.8. We then used cotyledons and the first real leaves from sterile tomato seedlings aged 1-2 weeks as explant material for transformation. A single colony of *A. tumefaciens* was cultured in LB medium supplemented with appropriate antibiotics overnight at 28°C. We resuspended the culture in inoculation buffer MMA (0.49% MS with vitamins + MES, 100µM Acetosyringone at pH 5.5-5.7) to an OD600 of 0.5, and inoculated the plant material with this for 20 minutes. We then co-cultivated the explants on GM infused with antibiotics at 22°C in the dark for two days. Antibiotics we used for selection included 20mg/L Hygromycin, 250mg/L Cefotaxime, and 150mg/L Timentin.

For callus/shoot regeneration from cotyledons, we followed the methodology described by (McCormick, 1991; Sun *et al*., 2006; Gupta and Van Eck, 2016; Sandhya *et al*., 2022). First, we selected for positive transformants on callus medium (CM; 1xMS with vitamins, MES, 3% sucrose, 1.5mg/L zeatin, 0.1mg/L IAA, and 0.8% plant agar, pH 5.8) containing suitable antibiotics in the dark at 21-26°C. Then, we transferred the developed callus with adventitious buds to shoot elongation medium (SM) in light which mirrored the CM in composition with the addition of 0.5-1mg/L zeatin. Then we transferred the developed shoots to rooting medium (RM) under long-day photoperiod to promote rooting.

### GUS staining assay

We characterized the promoter activity patterns with the first-generation transformed (T1) seeds. We grew the T1 plants two weeks in long day conditions and subsequently treated them with WL and WL+FR for 6 hours (ZT=3h to 9h). We assessed the activity of the β-glucuronidase (GUS) gene with a histochemical GUS staining assay. We carried out the staining with fresh staining solution (50mM NaPO4, 4mM potassium ferricyanide (III), 4mM potassium ferrocyanide (II), 0.05% (v/v) Triton X-100, 0.05% (w/v) X-Gluc). We submerged the seedlings in the freshly prepared staining solution and applied vacuum infiltration for 10-15 minutes to ensure thorough penetration. Following infiltration, the samples were incubated in darkness at 37°C for 24h when blue color started to appear. Plant tissue was washed with MQ water to remove excess stain, followed by a series of ethanol washes (50%, 70%, and 96%) for clearing which could be extended to an overnight duration if necessary. We then visualized the plants with microscopy and prepared sections as described above.

### Quantitative RT-PCR

To quantify changes to the transcript levels in response to supplemental FR, we used quantitative Real-Time (qRT) PCR. we subjected the plants to two different lighting conditions: exposure to white light (WL) and a low R/FR ratio of 0.2 (WL+FR). Post 6 or 24 hours of light treatment, we harvested the plant material, focusing on either the whole internode or the central cylinder hand dissected from the internode. We replicated this experiment four times, and for qPCR, we had three technical replicates. For each biological replicate, three internodes were collected into a tube, rapidly frozen in liquid nitrogen, and then stored at −80°C. We extracted total RNA using the RNeasy kit (Qiagen) and synthesised cDNA with random hexamer primers and RevertAid H Minus reverse transcriptase. We carried out real-time PCR with primers detailed in Supplementary Table S4 and SYBRgreen Super mix (Thermo Fisher Scientific) on the Bio-rad CFX Opus 384 system. To quantify the relative transcript abundance, we applied the comparative 2^ΔΔCt^ method, using *ACTIN2* as the reference gene for normalization (Reynoso *et al*., 2019). To statistically analyse the qPCR data we carried out ANOVA followed by a Tukey post hoc test using R software. For data presentation and visualization, we used GraphPad software (Version 9.5.1, released on January 24, 2023).

### Homolog analysis

In our study, we conducted a comprehensive homolog search across various plant species belonging to different families. We used the default BLAST search settings on several databases to source cDNA sequences. For Arabidopsis, we obtained sequences from TAIR (The Arabidopsis Information Resource, 2015). Soybean (*Glycine max*) and *Pisum sativum* sequences we sourced from Phytozome (Goodstein *et al*., 2012). Solanaceae family sequences we accessed from Sol Genomics (Solgenomics.net) (Mueller *et al*., 2005), which included ITAG4.0 for *Solanum lycopersicum*, *Capsicum annuum* UCD 10X genome chromosome v1.0 for *Capsicum annuum*, and eggplant V4 for *Solanum melongena*. Additionally, orthologs for *Brassica nigra* we obtained from the Brassica Database (http://brassicadb.cn/#/BLAST/).

In our methodology for identifying orthologs, we prioritized the highest score of similarity, maintaining a threshold of over 70%. In cases where the similarity difference was less than 10%, we included multiple homologs.

### Phylogenetic analyses

For our study, we aligned the homologs of all TFs using Muscle v3.8.31(Edgar, 2004). Following this alignment, we generated phylogenetic trees using the Maximum Likelihood method in the Mega X software (Kumar *et al*., 2016). This process was based on the Tamura Nei model for constructing the ML tree. To ensure the robustness of our phylogenetic analysis, we performed a bootstrap analysis with 1000 rapid bootstraps. Subsequently, we applied the consensus tree method to select representative genes for each species. We used the resulting output file from this process to construct a consensus tree following a majority rule consensus approach. This step was crucial to ensure that only nodes with bootstrap support values exceeding 50% were included in our final phylogenetic tree. We carried out the visual representation and fine-tuning of the ML analysis outcomes using Figtree v1.4.3 (Rambaut, 2012).

## Results

### Stem and pith cells elongated in response to FR in tomato

We characterized the tomato internode responses to low R:FR. To ensure our findings would be representative of tomato cultivars, we included tomatoes specifically bred for two different cultivation systems: M82 as a model for processing tomatoes and Moneymaker (MM) as a greenhouse-model. Our objective was to conduct detailed phenotyping on young plants to explore various aspects of stem responses at both the macroscopic and cellular levels. After transplanting germinated seedlings into soil and allowing a week for recovery, we subjected them to 7-day light treatments, either white light (WL) or white light supplemented with far-red (WL+FR). We gathered phenotypic data for both MM and M82 at 21 days after germination (dag), as shown in Figure 1a. We observed multiple phenotypic responses induced by WL+FR, with stem elongation being particularly prominent (Figure 1b). We chose the first internode for detailed cellular anatomy phenotyping, as it was the only internode present at the onset of FR treatment (14 days after sowing) and showed the strongest response. The cross and longitudinal sections were prepared at the internode’s midpoint, and a set of cell types (Figure 1c) were characterized for their cell layer numbers, areas, lengths, and widths. We discovered many cellular responses to FR (Figures 1d-e), such as Interfascicular Cambium thickness, Pith layer, Middle Pith length, Pith thickness, Parenchyma layer, Collenchyma layer, Collenchyma cell area, and Epidermis cell length (p<0.05 in WL vs WL+FR, Figure 1e). These findings provide a detailed understanding of the cellular changes in tomato plants in response to FR light treatments.

### Shade avoidance-related transcription factor identification by RNA-seq

We designed an RNA-seq experiment with multiple timepoints to identify the gene expression changes accompanying the internode elongation and the pith responses (Figure 2a). We selected timepoints that led to the discernible change in internode elongation (6h, based on Li *et al*. 2024). In addition to harvesting whole internode tissue, we were able to hand-dissect the central cylinder in plants of the latter two timepoints. These central cylinder samples are largely composed of pith but likely include some of the vascular tissues as well. We compared WL+FR samples at each timepoint to the corresponding WL samples to discern the FR-responsive differentially expressed genes (Figure 2b). Notably, at the 6-hour mark, the most DEGs were observed. Next, we tested the gene expression of the central cylinder against the whole internode for each treatment and timepoint (Figure 2c) and found a number of central cylinder-enriched genes. To delve deeper, we also intersected the upregulated DEGs between both the central cylinder and the internode at 30 and 48 hours, finding a higher count of unique DEGs in the central cylinder (Figure 2d). These are likely central cylinder-specific responses that are diluted out in the entire internode samples.

**Figure 2:**
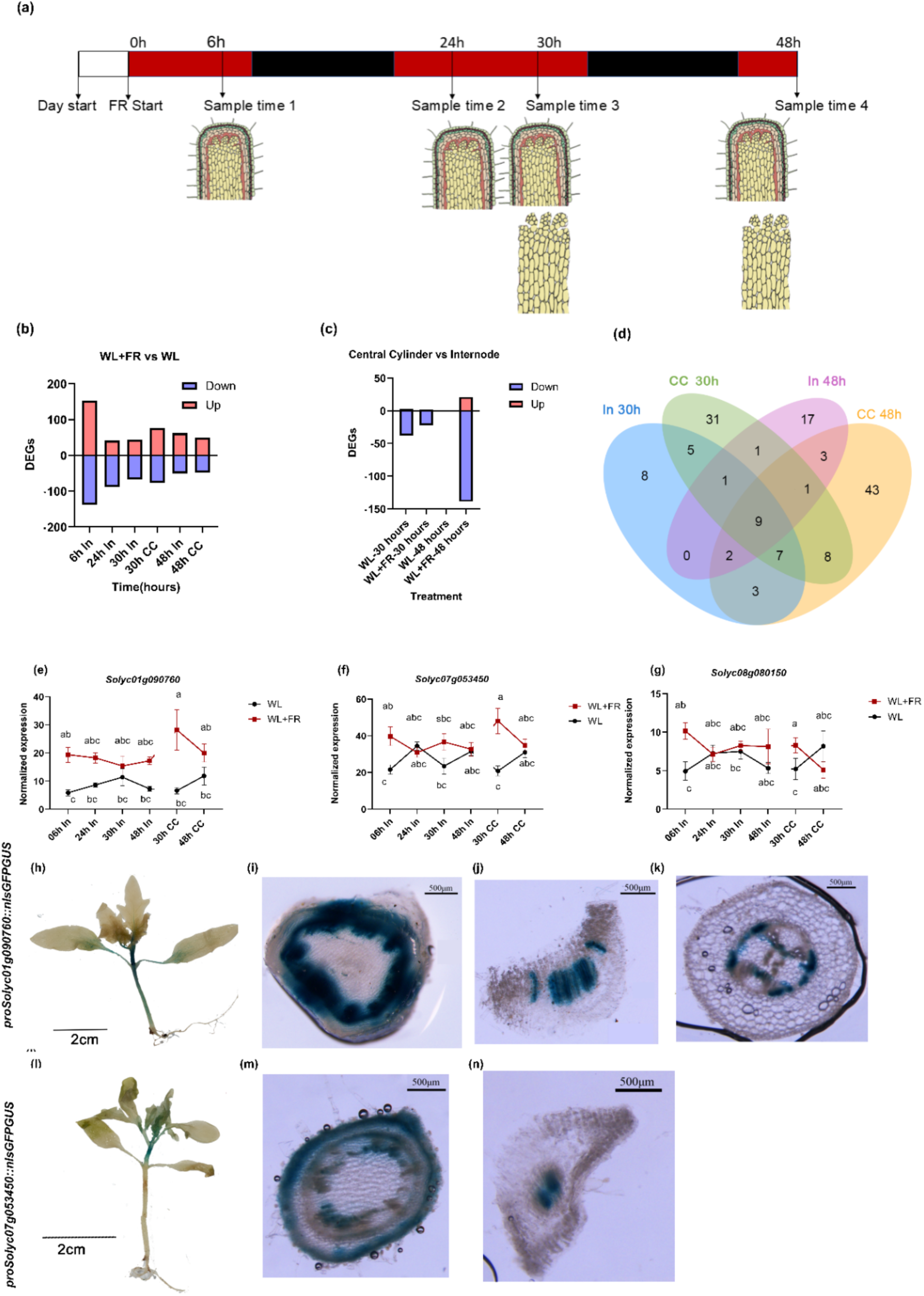
Transcriptomic responses of tomato first internode to low R:FR. a. The experimental design of the transcriptomics experiment, including the timing of harvest and the specific tissues harvested. White bar represents the WL treatment, red bars represent periods when the supplemental FR light is added, while black bars indicate night phases, and alternating light/dark bars depict the light/dark cycle. b. DEG analysis of FR-responsive genes for each timepoint for whole internode and central cylinder samples. Upregulated genes are marked in red and downregulated genes in blue. c. DEG analysis of central cylinder-enriched and -depleted genes, as identified by DE analysis for central cylinder vs whole internode for specific timepoints and light treatments. Central cylinder-enriched genes marked in red and central cylinder-depleted genes in blue. d. Overlap of FR-responsive upregulated DEGs between the central cylinder and whole internode at 30 h and 48 h of light treatment e-g. The expression patterns of the three genes encoding for TFs of interest. (e) *Solyc01g0S7C0*, (f) *Solyc07g05345S*, and (g) *Solyc08g080150*. The significant differences are determined using two-way ANOVA, and error bars represent Standard Errors h-k. GUS staining of 2-week-old *proSolyc01g0S07C0::nlsGFPGUS* seedling with 6h FR treatment. Cross sections show (i) the first internode, (j) petiole of the first true leaf and (k) the hypocotyl l-n. GUS staining of 2-week-old *proSolyc07g053450::nlsGFPGUS* seedling with 6h FR treatment. Cross sections show (m) the first internode and (n) petiole of the first true leaf.

We tested the FR-responsive central cylinder DEGs for GO term enrichment (Figure S2) to see if it resembled that of whole internode (Li *et al*., 2024), but we identified only a small number of GO terms. GO category “response to auxin” was enriched in the upregulated DEGs in the central cylinder as expected, but surprisingly only at the 30h timepoint and not at 48h. No other enriched GO terms were shared between FR-responsive DEGs of whole internode (Li *et al*., 2024) and central cylinder (Figure S2). We also tested GO term enrichment of the central cylinder-enriched DEGs (Figure S3) for each time point and treatment and GO category “transmembrane transport” was enriched throughout the central cylinder-enriched DEGs, either reflecting the little understood roles of pith, or the effect from the residual vascular tissues included in the sampling.

We further examined the DEG lists and identified all FR-responsive transcription factor (TF) encoding genes in our data (Table 1). TFs typically coordinate and amplify responses to environmental cues, so we focused our follow-up inquiry on selected TFs of interest. Specifically, we selected three upregulated TFs for detailed analysis based on their functions reported in the literature. These TFs, encoded by *Solyc08g080150*, *Solyc01g0S07C0*, and *Solyc07g053450*, showed upregulation in low R:FR in both the whole internode but also in the central cylinder (Figure 2e-g).

**Table 1.**
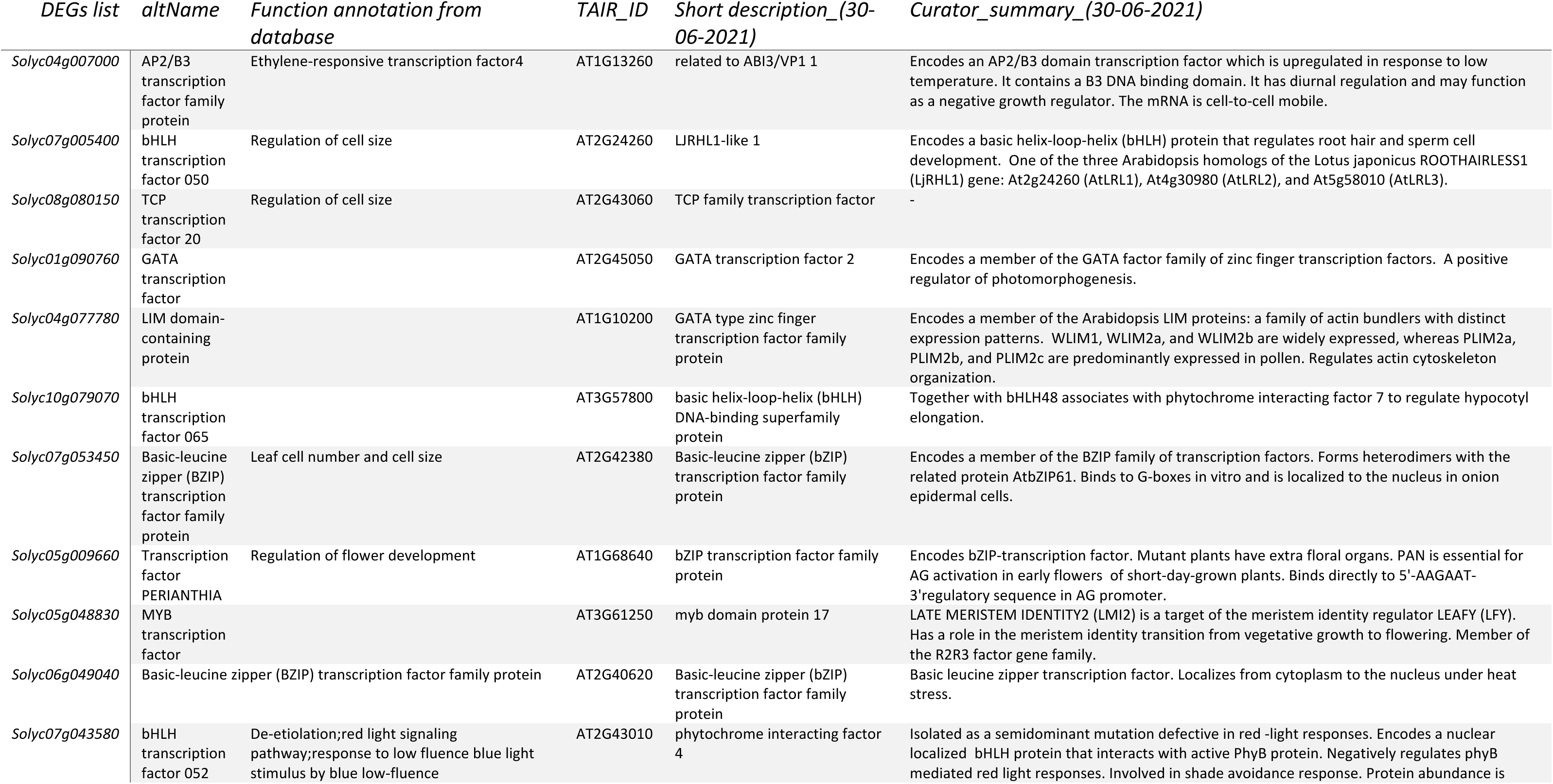
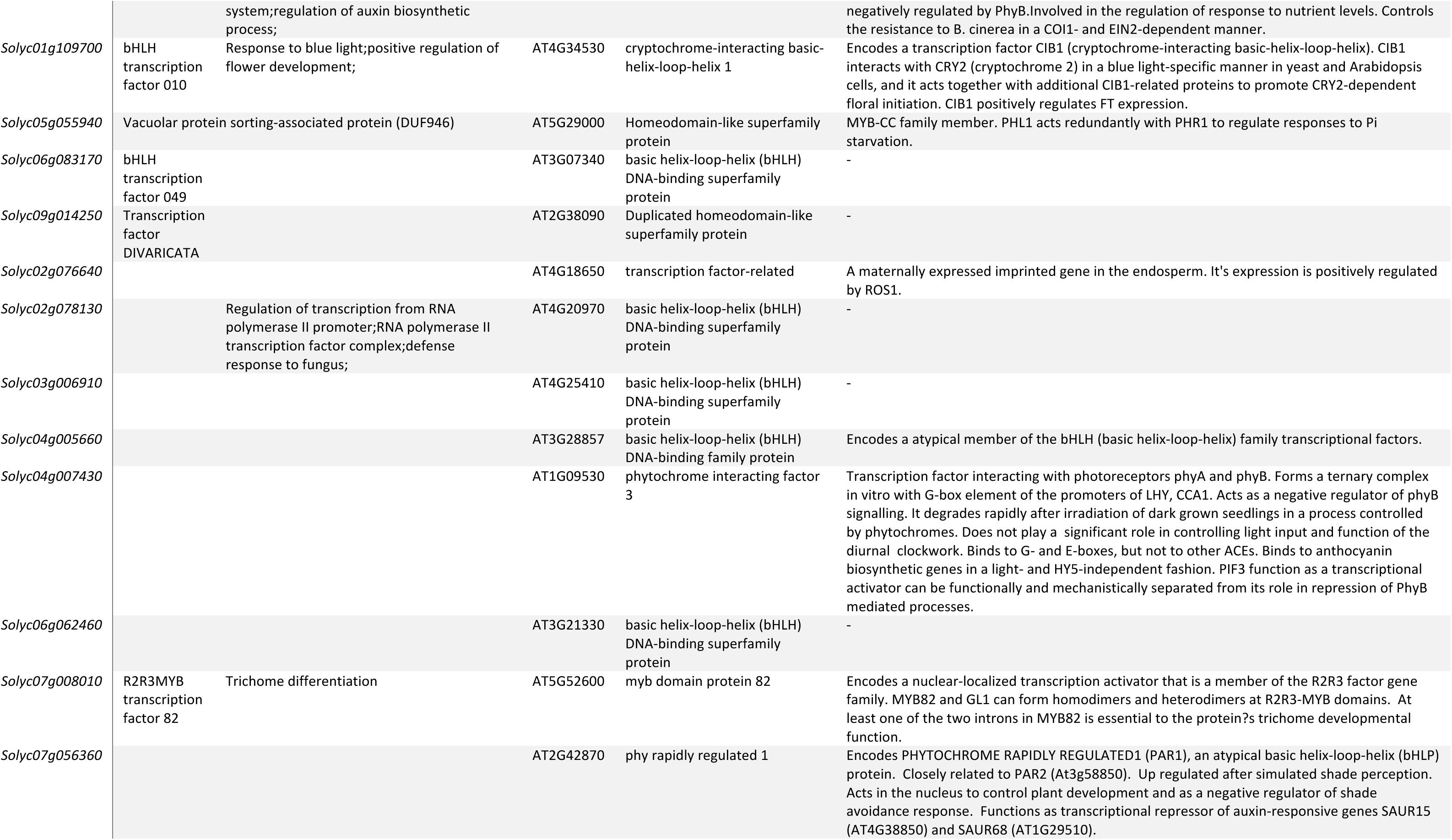
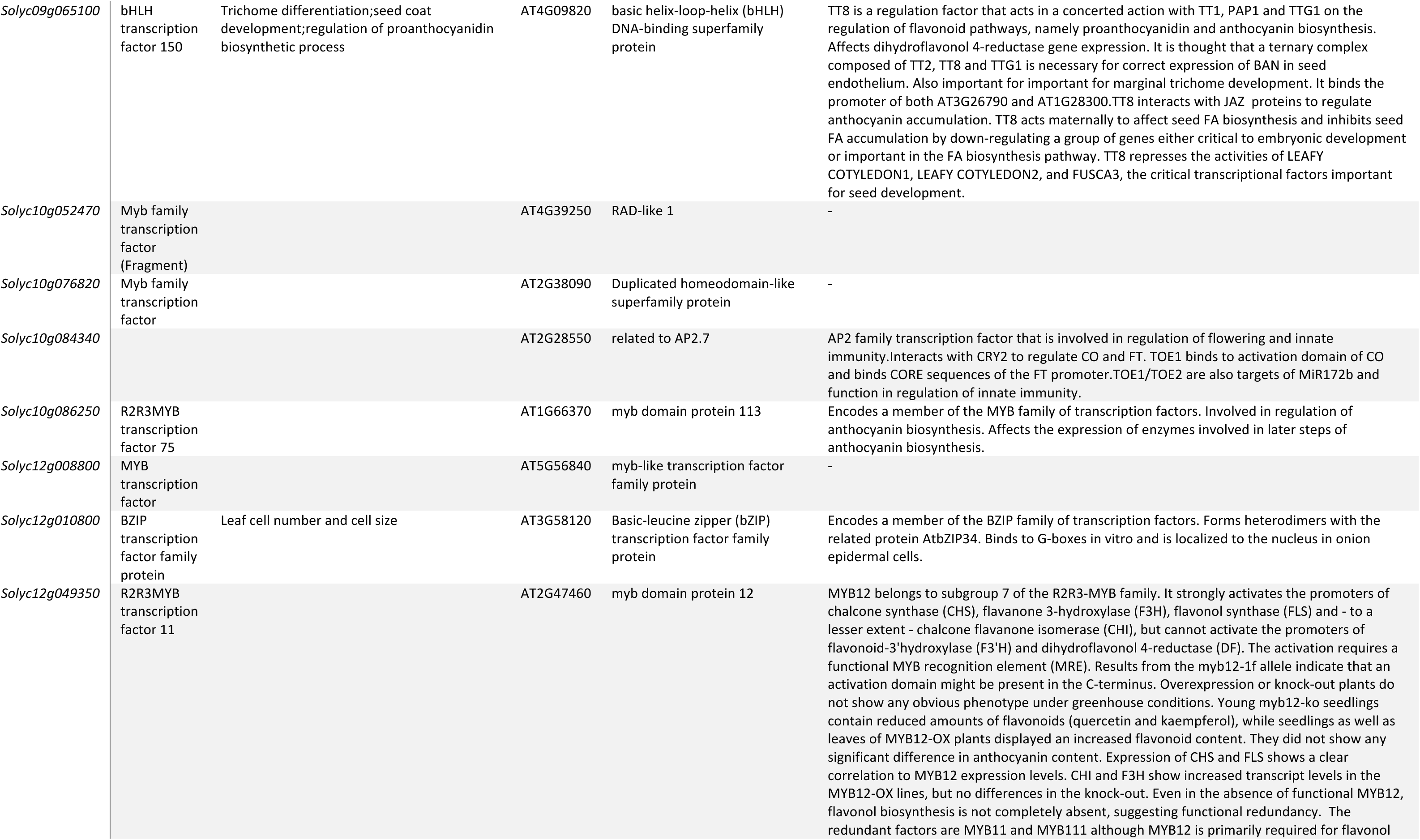
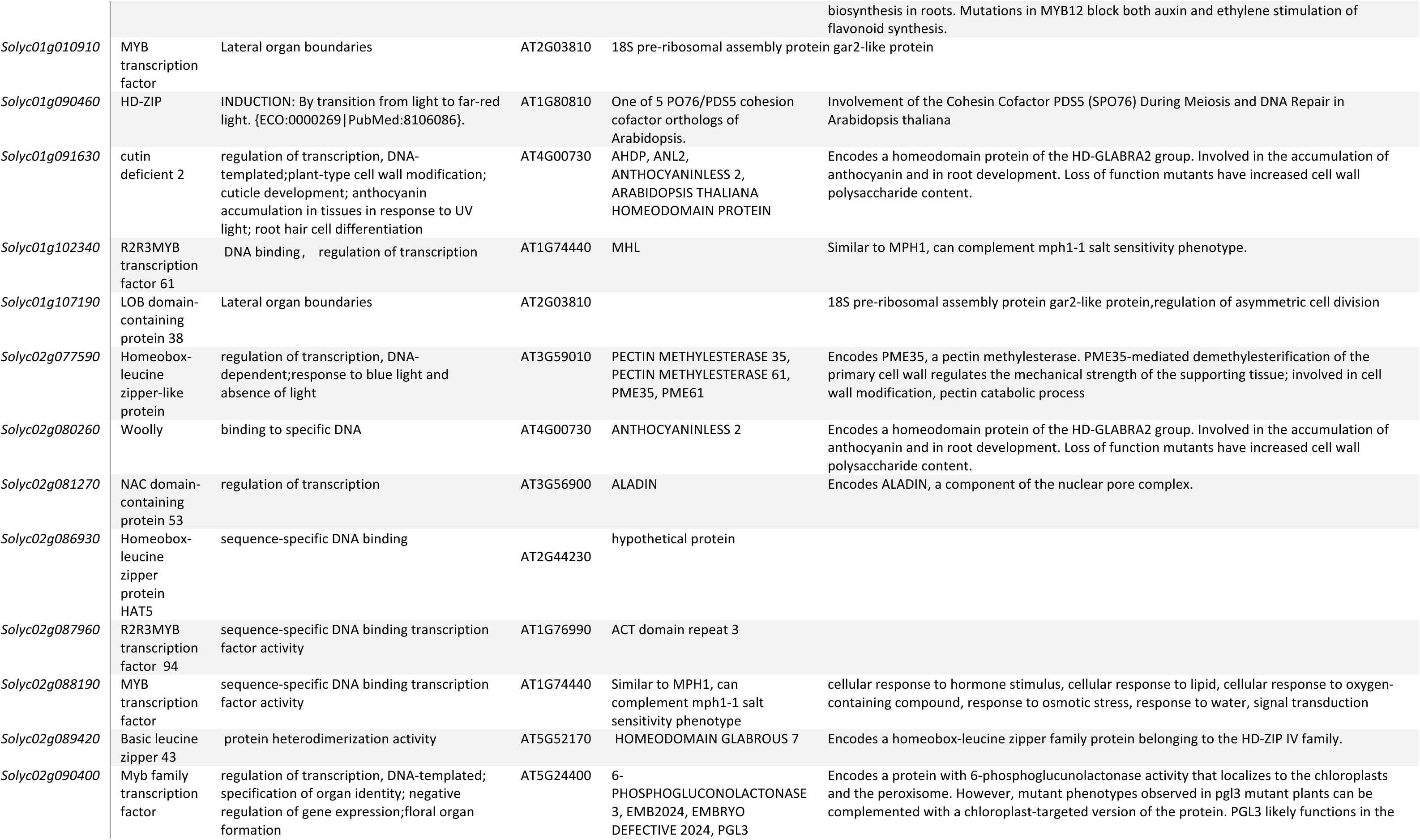
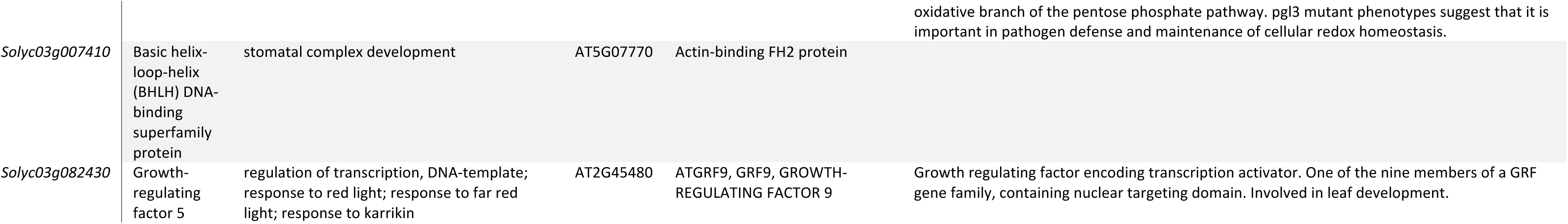
DEGs that are transcriptome factors. Functional annotations were obtained from Plant Transcription Factor Database (Jin et al., 2015; Jin et al., 2017), Solgenomics network (Fernandez-Pozo et al., 2015) and literature (Yang et al., 2015; Padmanabhan et al., 2022). Homologs from Arabidopsis were identified by multi-blast using Arabidopsis protein sequences.

*Solyc01g0S07C0* encodes for a GATA transcription factor from the zinc finger DNA binding protein family, with roles for development and cell differentiation (Gao *et al*., 2015). Its Arabidopsis homologs are implicated in longitudinal cell elongation, linking well with the stem and the cell elongation responses observed (Lu *et al*., 2021; Hou *et al*., 2022). *Solyc08g080150* encodes for SlTCP20, a TF with diverse roles in leaf development, flower symmetry regulation, shoot branching, and senescence involvement (Parapunova *et al*., 2014). Lastly, *Solyc07g053450* codes for a bZIP transcription factor, which is integral to several biological functions, notably in the response to various abiotic stresses (Liu *et al*., 2023*b*).

We wanted to test the spatial expression patterns driven by the promoters of these TFs. We constructed transcriptional reporter lines for *Solyc01g0S07C0* and *Solyc07g053450*, while we were not able to regenerate tomato lines with the *Solyc08g080150* transcriptional reporter construct. We visualized their expression with GUS staining, and this revealed distinct expression patterns for two TFs in tomato (Figure 2h-n). The *Solyc01g0S07C0* promoter showed abundant expression primarily in the hypocotyl, the first internode, and the petiole of true leaves. Detailed histological analysis narrowed down this expression to the xylem of the hypocotyl and first internode, as well as to young pith cells, and the central vascular bundle of the petiole. *Solyc07g053450* demonstrated specific expression in the internode and the petiole of true leaves, while conspicuously absent in the hypocotyl. Closer examination of cross-sections revealed its localization to the epidermis and young pith cells of the internode, and strictly to the stele, excluding the phloem, in the petiole. This differential expression suggests a role for Solyc07g053450 in regulating tissue-specific developmental processes, potentially influencing structural and physiological traits in these regions.

These findings provide insights into the spatial regulation of gene expression during early plant development, highlighting the intricate roles these TFs may play in tomato morphogenesis.

### Pith elongation is a conserved behavior in shade-avoiding species

After observing the cellular response to FR enrichment in tomato, we wondered if the pith phenotypes and specific transcriptional changes were limited to tomato, or if these responses were conserved across different dicot species with similar stem growth habits. We included two other Solanaceae family members, *Capsicum annuum* (bell pepper) and *Solanum melongena* (eggplant). For more distantly related species, we chose two pairs of species from the rosid clade. Firstly, we chose soybean (*Glycine max*) and *Pisum sativum* (pea), from the Fabaceae (legume) family and *Brassica nigra* (black mustard) and Arabidopsis from the Brassicaceae family. However, Arabidopsis does not form elongated internodes during juvenile growth, so there we looked into the inflorescence stem responses instead to test if this could serve as a model for stem responses. In these species, we observed a marked elongation response in internode 1 under supplemental FR light exposure (Figure 3). We also measured stem diameter and hypocotyl traits, and in some species, the elongation was accompanied by diameter increase (Figure S4). A common characteristic among these species, except for the Arabidopsis inflorescence, was the elongation of pith cells in the internode 1 (Figure 3). These findings establish tomato, bell pepper, eggplant, black mustard, and soybean as examples of FR-responsive species in our panel. Conversely, when testing pea, we noted no significant differences in stem length between the treatments (Figure 3), concluding that it is a species with lesser FR response. We also noted that the cellular arrangement of pea internodes was different to the other species we investigated (Figure S5). Cross-sections of pea stems revealed a tetrarch symmetry, creating an X-shaped pith, thus requiring specific directions for measurement (Figure S5). Here, to measure pith traits, we focused on the area predominantly composed of densely red-stained cells within the X-shaped area between vascular bundle. Our analysis showed no differences between treatments for the measured cell types in pea (Figure 3l), highlighting the variability in FR-responsiveness among different dicot species.

**Figure 3:**
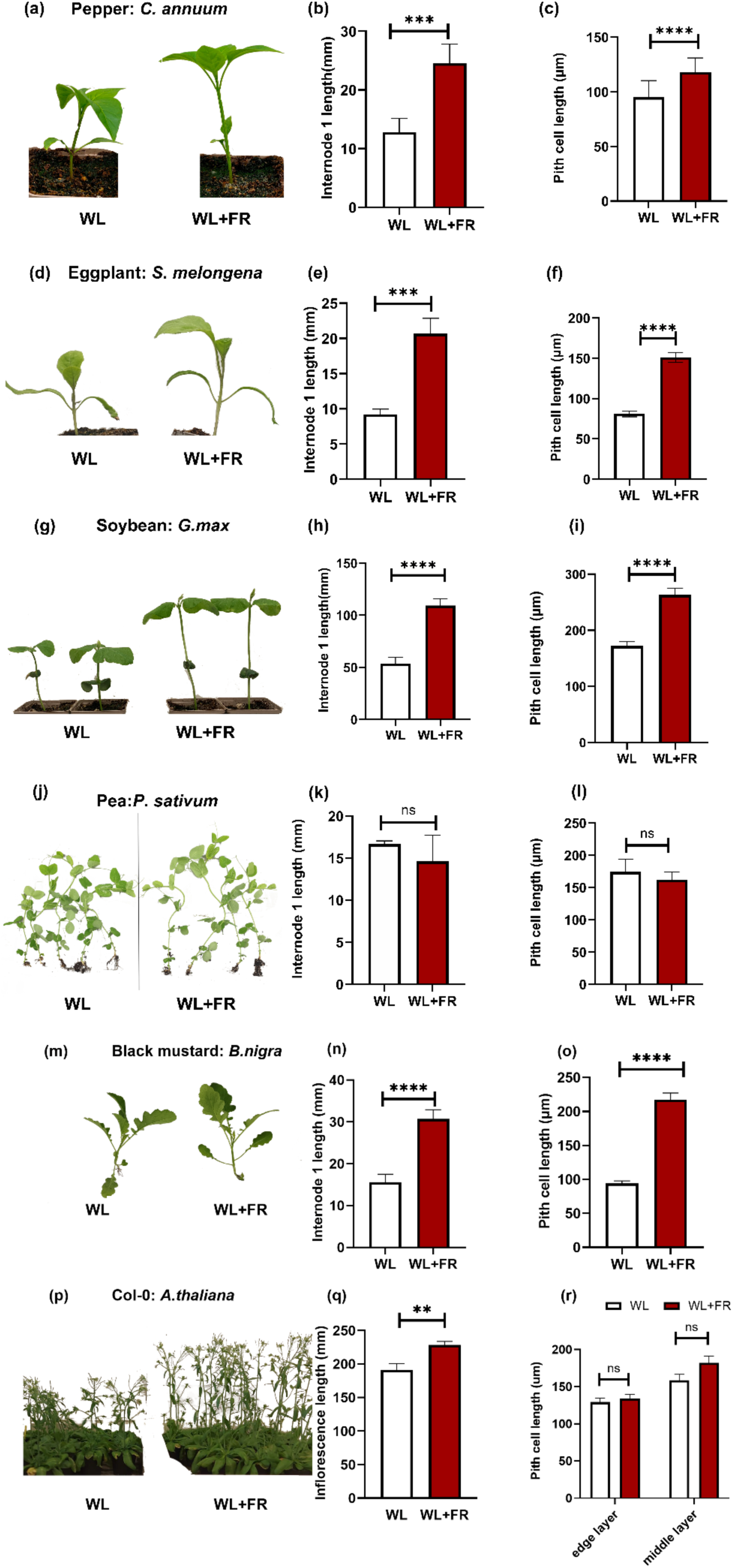
FR-responsive internode elongation is accompanied with pith cell changes across multiple dicots. Representative images of 7-day WL and WL+FR treated plants (a,d,g,j,m,p), measurements of the first internode or inflorescence length (b, e, h, k, n, w), and pith cell length measurements (c, f, i, l, n, r). The data is given by species as follows: (a-c) bell pepper (*Capsicum annuum*), (d-f) eggplant (*Solanum melongena*), (g-i) soybean (*Glycine max*), (j-l) pea (*Pisum sativum*), (m-o) black mustard (*Brassica nigra*), and (p-r) Arabidopsis (Col-0). Asterisks in the figure denote significant differences between WL and WL+FR conditions, with: * represents p≤ 0.05, ** p≤0.01, *** p≤0.001, and **** p≤0.0001. The sample size for each phenotyping comparison was approximately 12, and each experiment was repeated twice. The sample size for each cellular phenotyping comparison was approximately 50.

### FR-responsive tomato TFs have family-specific and tissue-specific expression patterns

Finally, we wanted to test if the molecular signatures coupled with internode and pith elongation responses to far-red (FR) light in tomatoes were conserved in our set of dicots. Hence, we tested if the FR-responsive behaviour of the transcripts for our candidate TFs identified in the tomato transcriptomics data, *Solyc07g053450, Solyc08g080150*, and *Solyc01g0S07C0*, was also observed in the other species.

For this analysis, we identified homologs for the three TF genes in all these species using BLAST (Fernandez-Pozo *et al*., 2015; The Arabidopsis Information Resource, 2015; Chen *et al*., 2022). In Solanaceae bell pepper and eggplant, we identified single hits for each of the three genes, whereas for Arabidopsis, homolog selection was guided by BLAST results and literature annotations to identify a homolog with a potentially conserved function. For pea, soybean, and black mustard, we chose genes based on high similarity scores (>80%). These homologs were visualized in maximum likelihood phylogenies (Figure 4), which generally reflected the relationships between the plant species and indicated recent gene or whole genome duplication events. To quantify the FR-responsive gene expression, we collected samples from whole first internodes and their central cylinder at 6 h and 24 h time points for WL and WL+FR light treatments. For pea, the central cylinder could not be dissected due to challenging morphology (Figure S5).

**Figure 4.**
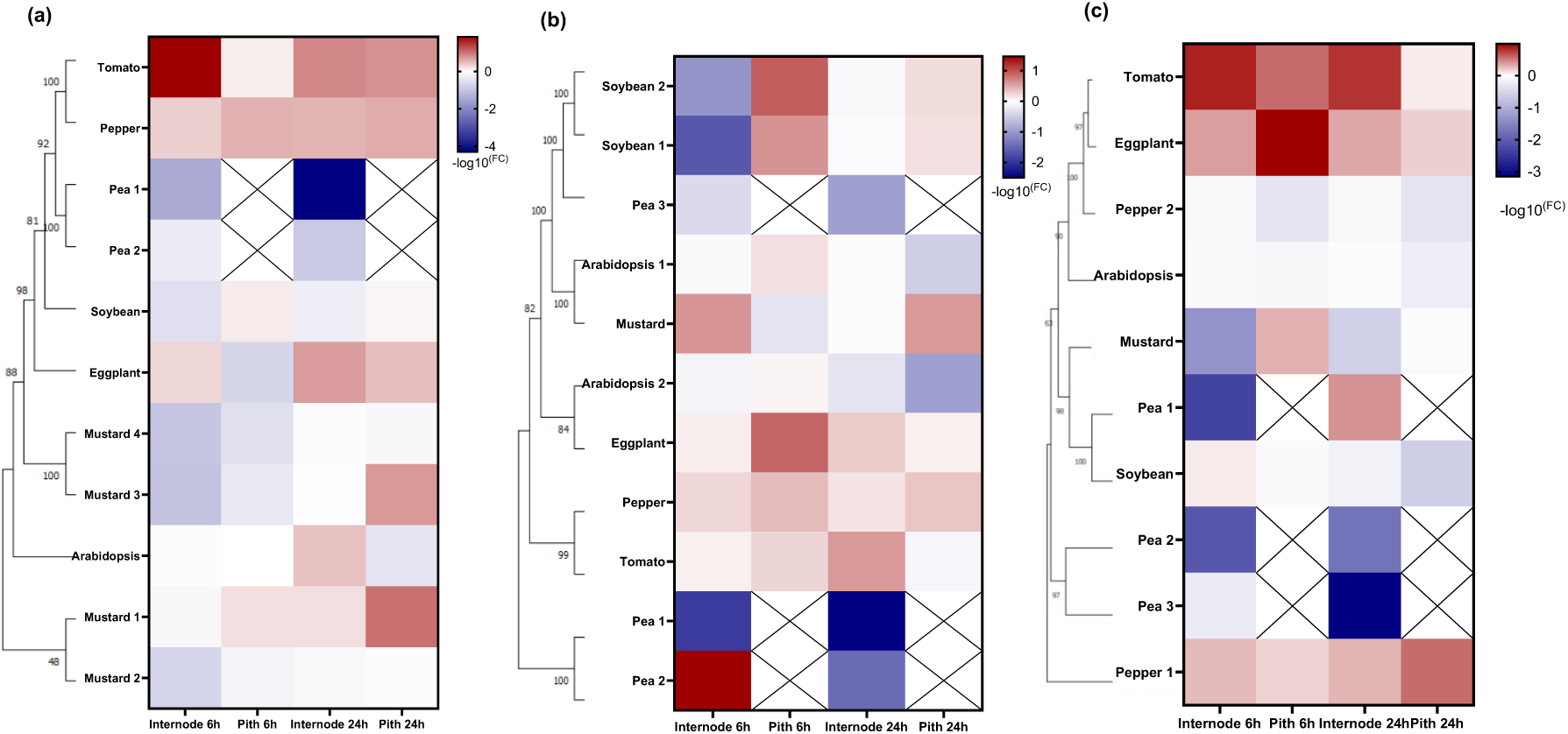
FR-responsiveness of target TF expression is conserved in Solanaceae. (a) Phylogeny and expression patterns of *Solyc07g053450* homologs: tomato (*Solyc07g053450*), bell pepper (*CA00g71840:1-921*), eggplant (*SMEL4.1_07g01S740.1.01*), Arabidopsis (*AT2G42380*), four black mustard homologs *(BniB06g001480.2N.1*, *BniB08g027470.2N.1*, *BniB08g072820.2N.1*, *BniB06g055220.2N.1*), soybean (*Gm0C:4S5378SS..4S53S1S5*), and two pea homologs (*XM_051027770.1*, *XM_05104545S.1*). (b) Phylogeny and expression patterns of *Solyc08g080150* homologs: tomato (*Solyc08g080150*), bell pepper (*TCP1S XM_01C7123S7.2*), eggplant (*chr8:85852001-85853900*), two Arabidopsis homologs (*AT2G45680*, *AT5G51910*), black mustard (*BniB08g029800.2N.1*), two soybean homologs (*Glyma.07G080300.1*, *Glyma07g08710.2*), and two pea homologs (*XM_0510304S0.1*, *XM_0510304S1.1*). (c) Phylogeny and expression patterns of *Solyc01g0S07C0* homologs: tomato (*Solyc01g0S07C0*), two bell pepper homologs (*chr3:2C7174801-2C717C100*, *chr8:35S02201-35S04000*), eggplant (*chr8:35S02201-35S04000*), Arabidopsis (*AT2G45050*), black mustard (*BniB07g039570.2N.1*), soybean (*GlysoPI4834C3.0CG078800.1*), and three pea homologs (*XM_051027377*, *XM_051030552.1*, *XM_051038772.1*). Each section utilizes Mega X software for phylogenetic analysis, adopting the Tamura Nei model with a bootstrap analysis (1000 replicates) for enhanced accuracy. The gene expression data from qRT-PCR is represented as heatmaps of log fold changes in gene expression under WL+FR versus WL conditions in the internode or pith at 6h and 24h after FR treatment, with red indicating upregulation, blue for downregulation, and white for no change (logFC=0).

The FR-responsive upregulation of *Solyc07g053450* and *Solyc08g080150* orthologs was consistent in both internode and central cylinder across the Solanaceae family: tomato, eggplant and bell pepper (Figure 4, S6, S7). For *Solyc01g0S07C0*, the conserved FR-induced upregulation was observed in tomato and eggplant, while in bell pepper, there were two homologs that showed divergent expression patterns. Unexpectedly, the bell pepper homolog that was less similar to *Solyc01g0S07C0* was FR-induced, while the homolog that was more similar had lost its FR-responsiveness (Figure 4, S8). This indicates that even with high sequence similarity, homologs can have diverged function, especially if another homolog has retained its role.

Outside the Solanaceae family, in Arabidopsis, black mustard, pea, and soybean, we generally observed a downregulation of these TF homologs in response to FR supplementation (Figure 4). In pea, where limited stem growth under FR was noted, all three TF homologs consistently showed decreased expression in response to FR. This suggests that these TFs might have acquired FR-responsive behavior within the Solanaceae lineage or lost it in the pea lineage. In soybean, which exhibited a less clear downregulation of the transcripts, the soybean homologs of *Solyc08g080150* showed strong FR-induction in the pith at 6h (Figure 4), indicating some conservation of function. In the FR-responsive soybean, while there is not such a clear downregulation in FR, there is also no clear homolog that would match the tomato patterns. For the two soybean homologs of *Solyc08g080150,* we observed a strong FR-induction specifically in central cylinder at 6h (Figure 4), so potentially the role of these two homologs is retained to some extent in soybean.

Thus, we observed that the expression patterns of FR-responsive genes were similar within the Solanaceae family but differ significantly from those in the Brassicaceae and Fabaceae families. Although certain species in Brassicaceae and Fabaceae showed elongation responses akin to those observed in Solanaceae, the expression patterns of the transcription factors (TFs) were not conserved across all families.

## Discussion

While extensive studies have been conducted on the role of the epidermis in SAS in Arabidopsis (Kohnen *et al*., 2016; Procko *et al*., 2016; Küpers *et al*., 2023), there is a notable gap in our understanding of the role of other cell types, and especially that of pith. The distinct anatomical and functional aspects of tomato stems and Arabidopsis petioles highlight this discrepancy. The pith, centrally located in the stem, is crucial for plant physiology and morphology but is often overlooked. This paper focuses on the pith responses and relates them to SAS across multiple species.

### FR-induced cell elongation, predominantly occurring in the pith in tomato, is likely conserved but with some exceptions

SAS has been extensively studied in Arabidopsis, where elongation traits in the hypocotyl and petiole show similarities to those seen in tomatoes (Chitwood *et al*., 2015). Research in Arabidopsis has highlighted a crucial aspect of SAS: the involvement of XTH (XYLOGLUCAN ENDOTRANSGLUCOSYLASE/HYDROLASE) proteins in cell wall relaxation, which aids cell expansion under shaded conditions (Sasidharan *et al*., 2010, 2014). This mechanism is instrumental in petiole elongation when exposed to supplemental FR light (Sasidharan *et al*., 2010). Specifically, the epidermis cells in the hypocotyl (18d) (Procko *et al*., 2016) and petiole (Pantazopoulou *et al*., 2017) of Arabidopsis become longer in response to supplemental FR light. In fact, the epidermis acts as a signal integrator in Arabidopsis and regulates organ elongation (Brown *et al*., 1995; Savaldi-Goldstein and Chory, 2008). However, the tissue morphologies and growth habits of Arabidopsis and tomato differ, making direct comparisons challenging. One of the differences between the cellular morphology of Arabidopsis petiole and tomato stem is the absence of pith cells in the Arabidopsis petiole. Pith cells, known as spongy parenchyma cells, provide support and nutrient storage in stems and roots. They are known for their extensibility, which differs from the outer layers (Abercrombie *et al*., 1968; Gallego-Giraldo *et al*., 2016; Yang *et al*., 2016). Additionally, pith undergoes programmed cell death as the tomato stem matures (Esau, 1953; Fujimoto *et al*., 2018). When we characterized the cellular anatomy responses of tomato cultivars M82 and Moneymaker to supplemental FR, we found a pronounced response of the pith to FR enrichment, with increased pith cell layers and cell length (Figure 1). While epidermis cells also became longer, their response was less pronounced compared to the pith. This led to a further investigation into how these cell types perceive and mediate SAS signals.

We profiled the FR-responsive gene expression patterns in the central cylinder of two tomato cultivars, M82 and Moneymaker (Figure 2). The central cylinder is largely composed of pith cells, but given the restrictions of hand dissection, some vascular cells are also likely to be present. We observed a notable increase in auxin-responsive genes, especially SAURs, under low R:FR conditions, with diurnal rhythm playing a key regulatory role (Li *et al*., 2024). The examination of central cylinder-specific transcriptomes revealed enrichment of “response to auxin” in the upregulated DEGs at 30 hours, but not at 48 hours. As the whole internode upregulated DEGs demonstrated enrichment of “response to auxin” that built up along the time course to 48 hours, this may be reflective of how auxin reaches the various tissues, likely inward out, and how the response builds up. It also may indicate that the auxin response is shorter lived in the central cylinder than in the outer tissue layers. We recently showed in Li *et al*., 2024 that the role of auxin in FR-responsive internode elongation in tomato is more complicated than in Arabidopsis, as exogenous Indole-3-acetic-acid (IAA) treatments were not able to recreate the full FR-induced elongation response, and we did not observe a FR-responsive increase in IAA in the internode. Taken together, this may support the importance of the tissue-specific distribution of auxin and its signaling. Additionally, the central cylinder demonstrated other significant differences from the whole internode, highlighting specific regulatory pathways (Figure S1). The involvement of various TFs in low R:FR response was evident, leading us to investigate these TFs further.

We then explored how conserved these responses were. We started with phenotyping of supplemental FR internode and pith responses in bell pepper, pea, soybean, eggplant, black mustard, and Arabidopsis (Figure 3). We observed that most species shared responses with tomato; FR-responsive internode elongation accompanied by increased pith cell length. Exceptions were pea with mild or no internode response and no cellular elongation (Ko *et al*., 2004; Skubisz *et al*., 2007). The conservation of this response in Arabidopsis and other plants is not well documented, but it is known that stem tissue expansion, including pith cells, is influenced by hormonal signals and genetic regulation (Nagata *et al*., 2004; Ye *et al*., 2021).

### Identification of tissue- and cell-type-specific FR-responsive TFs unique to Solanaceae

The identification of transcription factors (TFs) used in SAS is key to understanding how plants acclimate to variable light conditions. Uncovering these TFs helps understand the regulatory networks guiding the responses in shade-exposed plants. Additionally, recognizing tissue-specific expression patterns of these TFs adds power to the discovery, given that different plant tissues may respond uniquely to light variations. In our examination of FR-induced TF genes in tomatoes (Figure 2e-g), we focused on *Solyc01g0S07C0* (GATA TF), *Solyc08g080150* (TCP TF), and Solyc07g053450 (bZIP TF). We generated promoter reporter gene fusions of two of the TF-encoding genes, *Solyc01g0S07C0* and *Solyc07g053450*, revealing tissue- and cell-type-specific expression patterns in the internode central cylinder that extended into vasculature of petioles and leaves (Figure 2h-n).

Finally, we compared the FR-responsiveness of three tomato TF-encoding genes with their homologs in bell pepper, pea, soybean, eggplant, black mustard, and Arabidopsis (Figure 4). The observed divergence in FR-responsive expression patterns between the Solanaceae and the Brassicaceae/Fabaceae families raises compelling questions regarding the genetics and evolutionary aspects of the FR response. This variation leads to a hypothesis that the TF behaviors, characterized by upregulation in Solanaceae and downregulation in Brassicaceae/Fabaceae, might reflect a family-specific function induced by FR light in these TFs. This scenario opens new paths for investigating distinct regulatory mechanisms governing shade avoidance syndrome in the Solanaceae family, as opposed to other diverse plant families.

To contrast our data with that of existing Arabidopsis datasets, we utilized the eFP browser to examine Arabidopsis gene expression patterns of the TF homologs *AT5G51910, AT2G45050*, *AT2G45680*, and *AT2G42380* (Nakabayashi *et al*., 2005; Schmid *et al*., 2005; Winter *et al*., 2007). For the Arabidopsis homologs of *Solyc08g080150*, *AT5G51910*, exhibited increased expression in the inflorescence and seeds while *AT2G45680* showed high expression in leaves and the second internode of the inflorescence stem. For the *Solyc01g0S07C0* homolog *AT2G45050,* peak expression was in the petals. For *AT2G42380*, homolog of *Solyc07g053450*, demonstrated strong expression in mature pollen. These varied expression patterns indicate distinct roles for each homolog in specific developmental stages of Arabidopsis. Additionally, *AT2G45050*, *AT2G45680*, and *AT2G42380* have been reported to be upregulated in response to supplemental FR at the Arabidopsis leaf tip (Küpers *et al*., 2023). These genes were also found differentially expressed in the hypocotyl following supplemental FR seedling treatment (Kohnen *et al*., 2016), highlighting their FR-responsiveness in Arabidopsis tissues beyond those initially profiled in our study. This insight points towards a broader context of FR-responsive regulation in Arabidopsis that may extend beyond the developmental stages and tissues we initially focused on.

The lack of conservation in TF expression across different species suggests that the regulatory pathways activating these TFs have undergone evolutionary divergence, leading to distinct gene expression patterns. Conservation of TF binding events often correlates with the conservation of gene expression at the gene level (Hemberg and Kreiman, 2011). However, this conservation can vary significantly across species and tissues (Diehl and Boyle, 2018). Thus, the disparity in TF expression across different species implies that the regulatory mechanisms governing these TFs have evolved uniquely, potentially resulting in species-specific or tissue-specific expression patterns. This finding reminds us of the complex nature of evolutionary adaptations in plant responses and importance in characterizing traits in different plant families.

### Perspectives on the role of pith in SAS

Overall, we observed concurrent internode and pith elongation across the species studied, with transcriptionally typically inactive pith showing increased expression of these TFs in Solanaceae. However, our data do not conclusively link the pith function to shade avoidance elongation, and this remains to be firmly established.

Future research directions include extending studies to closely related families of Solanaceae and beyond, to pinpoint when FR-responsive behavior of these TFs ceases. Functional validation of these TF genes in shade avoidance through knockout and induction experiments would clarify their role. Additionally, investigating plants with varied pith morphologies could offer further insight. Specifically, it would be intriguing to determine if FR-responsive pith elongation and internode elongation are decoupled in any dicot species, shedding light on the diversity of plant acclimation strategies to light environments.

## Data availability

The RNA sequencing data from this study are openly available in NCBI GEO repository reference number GSE255611.

## Acknowledgments

We want to express our gratitude to Dr. Peter Etchells feedback and guidance and Ms. Annemieke Nieuwhof for technical support in this work. This work was supported by China Scholarship Council (CSC) PhD fellowship to LL. KK is supported by NWO Vidi grant number VI.Vidi.193.104, and RP is supported by NWO Vici grant number 865.17.002

## Contributions

KK, RP: Conceptualization and Supervision.

LL: Funding Acquisition, Data Curation, Formal Analysis, Visualization, Writing – Original Draft.

LL, YK, LvdK: Investigation.

LL, KK, RP: Writing – Review C Editing.

All authors read and have approved the final manuscript.

## Conflict of Interest

Authors declare no conflict of interest.

## Supplementary files

Table S1: DEGs (limma-voom output) – Both FR-responsive DEGs AND pith-specific DEGs

Table S2: GO enrichment analysis – FR responsive and pith-specific.

Table S3: Sequences of the synthesized promoters. Table S4: qPCR primers used in this study.

Figure S1: The number of differentially expressed genes (DEGs) between the Moneymaker and M82 cultivars in tomato internodes at each sampled timepoint and treatment.

Figure S2: GO enrichment analysis of FR-responsive (WL+FR vs WL) DEG in the central cylinder of each cultivar.

Figure S3: GO enrichment analysis of central cylinder-specific DEGs, focusing on the comparison between the central cylinder (CC) and the entire internode.

Figure S4: Shoot traits measured in multiple dicot species under WL and WL+FR treatments.

Figure S5: Cross section of pea (P. sativum) internode and cell type identification.

Figure S6: Fold change of transcript abundance of Solyc07g053450 homologs in response to FR treatment.

Figure S7: Fold change of transcript abundance of Solyc08g080150 homologs in response to FR treatment.

Figure S8: Fold change of transcript abundance of Solyc01g090760 homologs in response to FR treatment.

## Notes

### Competing Interest Statement

The authors have declared no competing interest.

## References

Abercrombie M (Michael), Brachet J (Jean), King TJ. 1968. Advances in morphogenesis. Volume 7. Elsevier Science.

Avalos G, Avalos G. 2019. Shade tolerance within the context of the successional process in tropical rain forests. Revista de Biología Tropical 67, 53–77.

Ballaré CL, Pierik R. 2017. The shade-avoidance syndrome: Multiple signals and ecological consequences. Plant Cell and Environment 40, 2530–2543.

Brown CL, Sommer HE, Pienaar L V. 1995. The predominant role of the pith in the growth and development of internodes in Liquidambar styraciflua (Hamamelidaceae). I. Histological basis of compressive and tensile stresses in developing primary tissues. American Journal of Botany 82, 769–776.

Chen H, Wang T, He X, Cai X, Lin R, Liang J, Wu J, King G, Wang X. 2022. BRAD V3.0: an upgraded Brassicaceae database. Nucleic Acids Research 50, D1432–D1441.

Chitwood DH, Kumar R, Ranjan A, et al. 2015. Light-Induced Indeterminacy Alters Shade-Avoiding Tomato Leaf Morphology. Plant Physiology 169, 2030–2047.

Coverdale TC, Agrawal AA. 2021. Evolution of shade tolerance is associated with attenuation of shade avoidance and reduced phenotypic plasticity in North American milkweeds. American Journal of Botany 108, 1705–1715.

Diehl AG, Boyle AP. 2018. Conserved and species-specific transcription factor co-binding patterns drive divergent gene regulation in human and mouse. Nucleic Acids Research 46, 1878–1894.

Edgar RC. 2004. MUSCLE: A multiple sequence alignment method with reduced time and space complexity. BMC Bioinformatics 5, 113.

Esau K. 1953. Plant anatomy. *in press.*

Fernandez-Pozo N, Menda N, Edwards JD, et al. 2015. The Sol Genomics Network (SGN)--from genotype to phenotype to breeding. Nucleic acids research 43, D1036–D1041.

Fujimoto M, Sazuka T, Oda Y, et al. 2018. Transcriptional switch for programmed cell death in pith parenchyma of sorghum stems. Proceedings of the National Academy of Sciences of the United States of America 115, E8783–E8792.

Gallego-Giraldo L, Shadle G, Shen H, Barros-Rios J, Fresquet Corrales S, Wang H, Dixon RA. 2016. Combining enhanced biomass density with reduced lignin level for improved forage quality. Plant Biotechnology Journal 14, 895–904.

Gao J, Chen YH, Peterson LAC. 2015. GATA family transcriptional factors: Emerging suspects in hematologic disorders. Experimental Hematology and Oncology 4, 1–7.

van Gelderen K, Kang C, Paalman R, Keuskamp D, Hayes S, Pierik R. 2018. Far-Red Light Detection in the Shoot Regulates Lateral Root Development through the HY5 Transcription Factor. The Plant Cell 30, 101–116.

Gommers CMM, Keuskamp DH, Buti S, van Veen H, Koevoets IT, Reinen E, Voesenek LACJ, Pierik R. 2017. Molecular profiles of contrasting shade response strategies in wild plants: Differential control of immunity and shoot elongation. Plant Cell 29, 331–344.

Goodstein DM, Shu S, Howson R, et al. 2012. Phytozome: a comparative platform for green plant genomics. Nucleic Acids Research 40, D1178–86.

Gupta S, Van Eck J. 2016. Modification of plant regeneration medium decreases the time for recovery of Solanum lycopersicum cultivar M82 stable transgenic lines. Plant Cell, Tissue and Organ Culture 127, 417–423.

Hagen G, Guilfoyle T, Weinig C, et al. 2015. Plasticity versus canalization: Population differences in the timing of shade-avoidance responses. Frontiers in Plant Science 54, 1–11.

Hemberg M, Kreiman G. 2011. Conservation of transcription factor binding events predicts gene expression across species. Nucleic Acids Research 39, 7092.

Hou Q, Zhao W, Lu L, et al. 2022. Overexpression of HLH4 Inhibits Cell Elongation and Anthocyanin Biosynthesis in Arabidopsis thaliana. Cells, 1–18.

Ko JH, Han KH, Park S, Yang J. 2004. Plant body weight-induced secondary growth in Arabidopsis and its transcription phenotype revealed by whole-transcriptome profiling. Plant Physiology 135, 1069–1083.

Kohnen M V., Schmid-Siegert E, Trevisan M, et al. 2016. Neighbor detection induces organ-specific transcriptomes, revealing patterns underlying hypocotyl-specific growth. Plant Cell 28, 2889–2904.

Kumar S, Stecher G, Tamura K. 2016. MEGA7: Molecular Evolutionary Genetics Analysis Version 7.0 for Bigger Datasets. Molecular Biology and Evolution 33, 1870–1874.

Küpers JJ, Snoek BL, Oskam L, et al. 2023. Local light signaling at the leaf tip drives remote differential petiole growth through auxin-gibberellin dynamics. Current Biology 33, 75–85.e5.

Li L, Wonder J, Helming T, Asselt G van, Pantazopoulou CK, Kaa Y van de, Kohlen W, Pierik R, Kajala K. 2024. Evaluation of the roles of brassinosteroid, gibberellin and auxin for tomato internode elongation in response to low red:far-red light. Physiologia Plantarum 176, e14558.

Liu X, Joseph F. Miceli I, Patton S, Murray M, Evans J, Wei X, Wang P. 2023a. Agrobacterial Transformation Enhancement by Improved Competent Cell Preparation and Optimized Electroporation. Life 13, 2217.

Liu H, Tang X, Zhang N, Li S, Si H. 2023b. Role of bZIP Transcription Factors in Plant Salt Stress. International Journal of Molecular Sciences 24.

Lu R, Zhang J, Wu YW, Wang Y, Zhang J, Zheng Y, Li Y, Li XB. 2021. bHLH transcription factors LP1 and LP2 regulate longitudinal cell elongation. Plant Physiology 187, 2577–2591.

Mathews S. 2006. Phytochrome-mediated development in land plants: red light sensing evolves to meet the challenges of changing light environments. Molecular Ecology 15, 3483–3503.

McCormick S. 1991. Transformation of tomato with Agrobacterium tumefaciens. Plant Tissue Culture Manual, 311–319.

Miyazawa Y, Manythong C, Fukuda S, Ogata K. 2014. Comparison of the growth traits of a commercial pioneer tree species, paper mulberry (Broussonetia papyrifera L. vent.), with those of shade-tolerant tree species: Investigation of the ecophysiological mechanisms underlying shade-intolerance. Agroforestry Systems 88, 907–919.

Mueller LA, Solow TH, Taylor N, et al. 2005. The SOL Genomics Network. A comparative resource for Solanaceae biology and beyond. Plant Physiology 138, 1310–1317.

Nagata T, Todoriki S, Kikuchi S. 2004. Radial expansion of root cells and elongation of root hairs of Arabidopsis thaliana induced by massive doses of gamma irradiation. Plant and Cell Physiology 45, 1557–1565.

Nakabayashi K, Okamoto M, Koshiba T, Kamiya Y, Nambara E. 2005. Genome-wide profiling of stored mRNA in Arabidopsis thaliana seed germination: epigenetic and genetic regulation of transcription in seed. The Plant journal : for cell and molecular biology 41, 697–709.

Osborne DJ. 1991. Light and plant responses. A study of plant photophysiology and the natural environment. Endeavour 15, 37.

Pantazopoulou CK, Bongers FJ, Küpers JJ, Reinen E, Das D, Evers JB, Anten NPRR, Pierik R. 2017. Neighbor detection at the leaf tip adaptively regulates upward leaf movement through spatial auxin dynamics. Proceedings of the National Academy of Sciences 114, 7450–7455.

Parapunova V, Busscher M, Busscher-Lange J, Lammers M, Karlova R, Bovy AG, Angenent GC, De Maagd RA. 2014. Identification, cloning and characterization of the tomato TCP transcription factor family. BMC Plant Biology 14, 1–17.

Pierik R, De Wit M. 2014. Shade avoidance: Phytochrome signalling and other aboveground neighbour detection cues. Oxford University Press.

Procko C, Burko Y, Jaillais Y, Ljung K, Long JA, Chory J. 2016. The epidermis coordinates auxin-induced stem growth in response to shade. Genes and Development 30, 1529–1541.

Rambaut A. 2012. FigTree v1. 4. and epidemiology. Edinburgh, UK: University of Edinburgh *in press*.

Reynoso MA, Kajala K, Bajic M, et al. 2019. Evolutionary flexibility in flooding response circuitry in angiosperms. Science 365, 1291–1295.

Ron M, Kajala K, Pauluzzi G, et al. 2014. Hairy root transformation using Agrobacterium rhizogenes as a tool for exploring cell type-specific gene expression and function using tomato as a model Mily. Plant Physiology 166, 455–469.

Sandhya D, Jogam P, Venkatapuram AK, Savitikadi P, Peddaboina V, Allini VR, Abbagani S. 2022. Highly efficient Agrobacterium-mediated transformation and plant regeneration system for genome engineering in tomato. Saudi Journal of Biological Sciences 29, 103292.

Sasidharan R, Chinnappa CC, Staal M, Elzenga JTM, Yokoyama R, Nishitani K, Voesenek LACJ, Pierik R. 2010. Light quality-mediated petiole elongation in Arabidopsis during shade avoidance involves cell wall modification by xyloglucan endotransglucosylase/hydrolases. Plant Physiology 154, 978–990.

Sasidharan R, Keuskamp DH, Kooke R, Voesenek LACJCJ, Pierik R. 2014. Interactions between Auxin, Microtubules and XTHs Mediate Green Shade-Induced Petiole Elongation in Arabidopsis. (E Huq, Ed.). PLoS ONE G, e90587.

Savaldi-Goldstein S, Chory J. 2008. Growth coordination and the shoot epidermis.

Schmid M, Davison TS, Henz SR, Pape UJ, Demar M, Vingron M, Schölkopf B, Weigel D, Lohmann JU. 2005. A gene expression map of Arabidopsis thaliana development. 37, 501–506.

Skubisz G, Kravtsova TI, Velikanov LP. 2007. Analysis of the strength properties of pea stems. International Agrophysics 21, 189–197.

Smith H, Whitelam GC. 1997. The shade avoidance syndrome: Multiple responses mediated by multiple phytochromes. Plant, Cell and Environment 20, 840–844.

Sottosanti K. 2023. Pioneer species | Definition, Examples, Ecology, & Facts | Britannica. https://www.britannica.com/science/pioneer-species. Accessed November 2023.

Sun HJ, Uchii S, Watanabe S, Ezura H. 2006. A highly efficient transformation protocol for Micro-Tom, a model cultivar for tomato functional genomics. Plant & cell physiology 47, 426–431.

The Arabidopsis Information Resource. 2015. TAIR -Home Page. https://www.arabidopsis.org/index.jsp. Accessed July 2022.

Tomescu AMF, Mcqueen CR. 2022. A protoxylem pathway to evolution of pith? An hypothesis based on the Early Devonian euphyllophyte Leptocentroxyla. Annals of Botany 130, 785–798.

Winter D, Vinegar B, Nahal H, Ammar R, Wilson G V., Provart NJ. 2007. An ‘Electronic Fluorescent Pictograph’ browser for exploring and analyzing large-scale biological data sets. PloS one 2.

de Wit M, Kegge W, Evers JB, Vergeer-van Eijk MH, Gankema P, Voesenek LACJ, Pierik R. 2012. Plant neighbor detection through touching leaf tips precedes phytochrome signals. Proceedings of the National Academy of Sciences of the United States of America 109, 14705–10.

Yang L, Zhao X, Yang F, Fan D, Jiang Y, Luo K. 2016. PtrWRKY19, a novel WRKY transcription factor, contributes to the regulation of pith secondary wall formation in Populus trichocarpa. Scientific Reports 6, 1–12.

Ye L, Wang X, Lyu M, Siligato R, Eswaran G, Vainio L, Blomster T, Zhang J, Mähönen AP. 2021. Cytokinins initiate secondary growth in the Arabidopsis root through a set of LBD genes. Current Biology 31, 3365–3373.e7.

Zabel RA (Robert A., Morrell JJ. 2020. Wood microbiology : decay and its prevention. Academic Press.

