## Supplementary figures S1-S7 for "Pith cell responses to low red: far-red light in dicot stems"

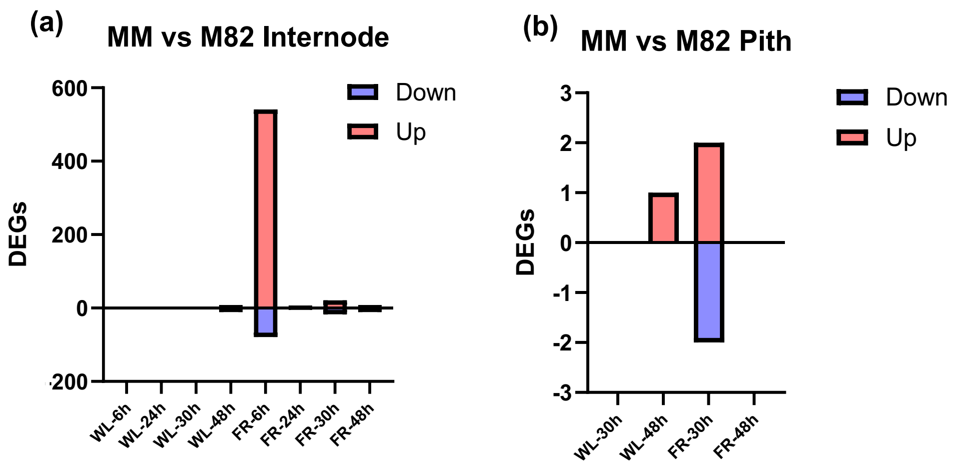


**Figure S1: The number of differentially expressed genes (DEGs) between the Moneymaker and M82 cultivars in tomato internodes at each sampled timepoint and treatment.** (a) Internode DEGs in WL or WL+FR treatment. (b) Central cylinder DEGs in WL or WL+FR treatment.

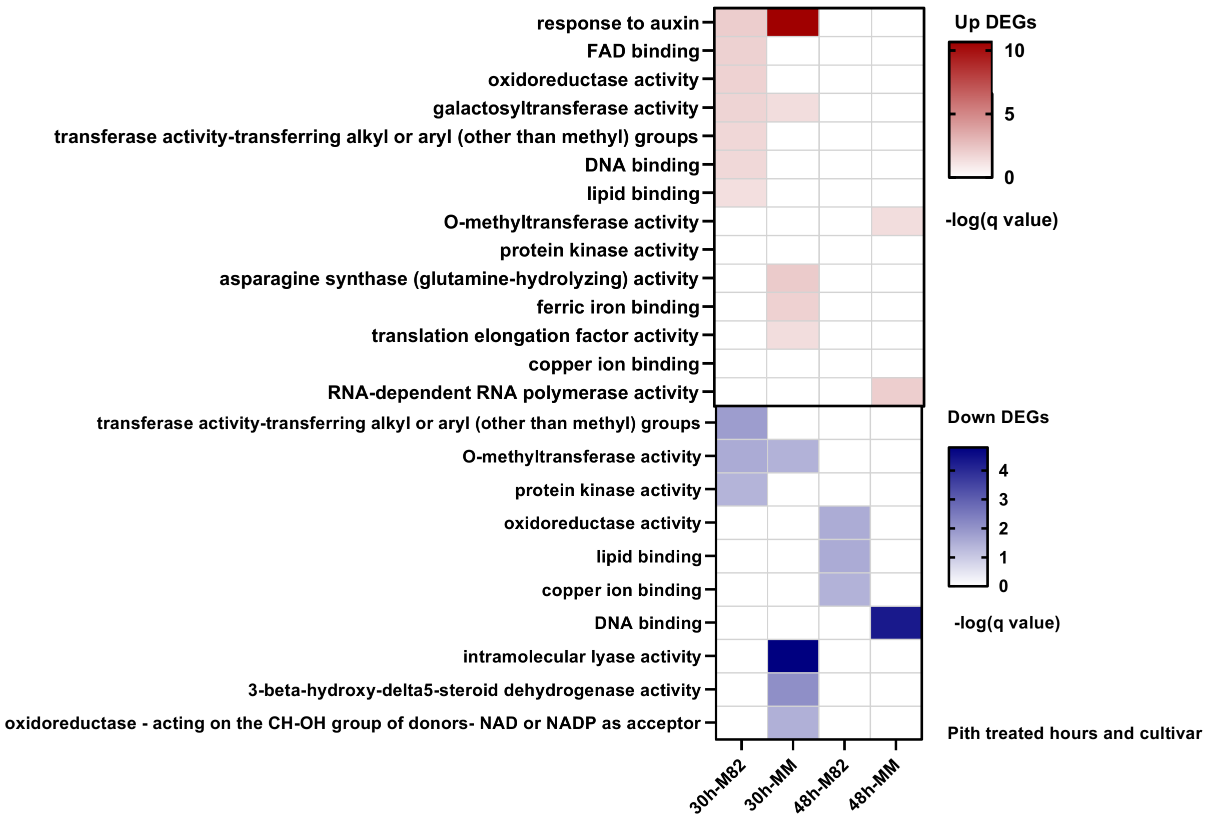


**Figure S2: GO enrichment analysis of FR-responsive (WL+FR vs WL) DEG in the central cylinder of each cultivar.** The heatmap displays the negative logarithm of the adjusted p-values (q-values) for each GO category, using a red color scheme to represent upregulated DEGs and blue for downregulated ones.

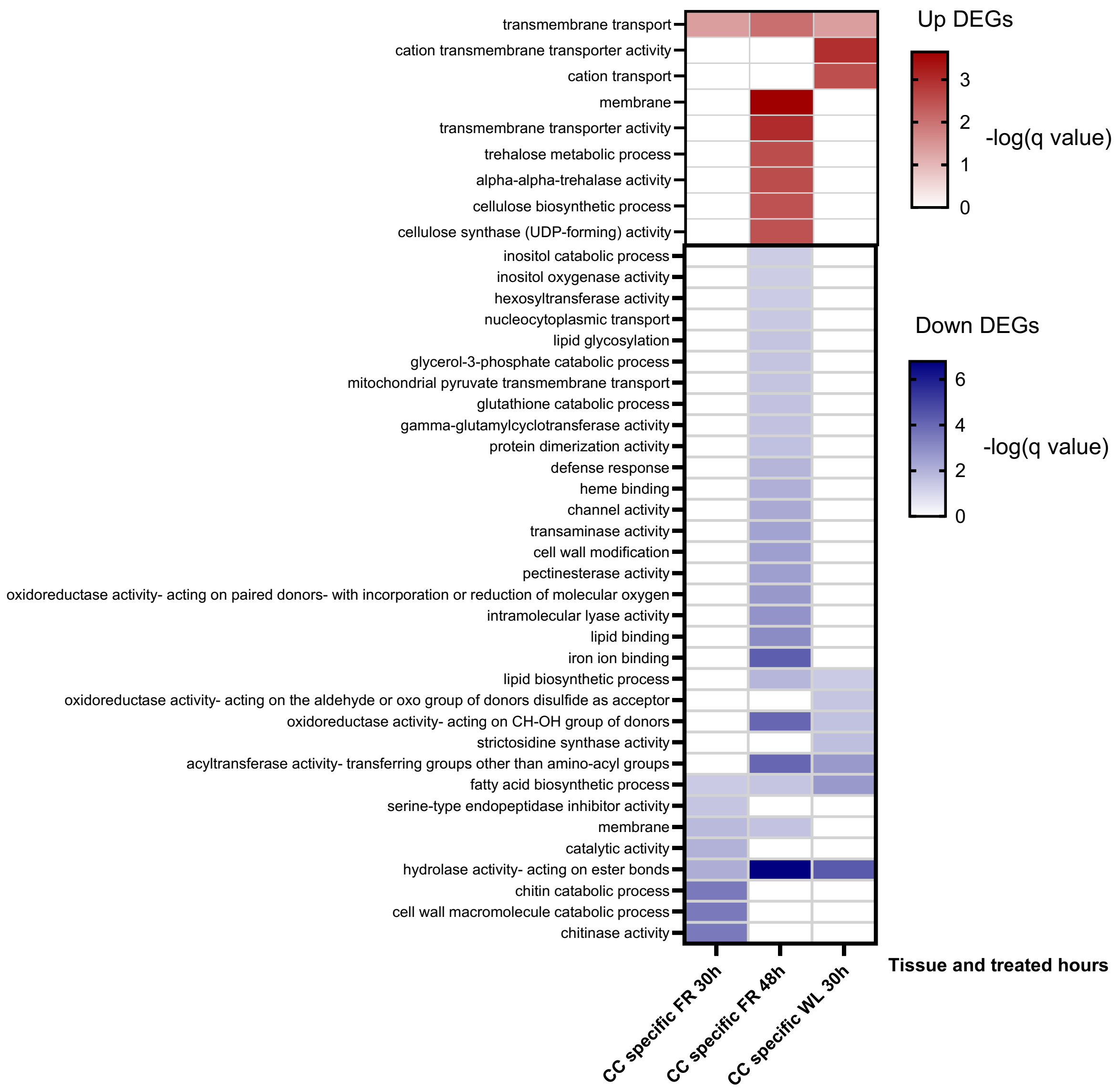


**Figure S3: GO enrichment analysis of central cylinder-specific DEGs, focusing on the comparison between the central cylinder (CC) and the entire internode.** The heatmap details the -log of adjusted q-values for each enriched GO category, with red indicating upregulated DEGs and blue for downregulated.

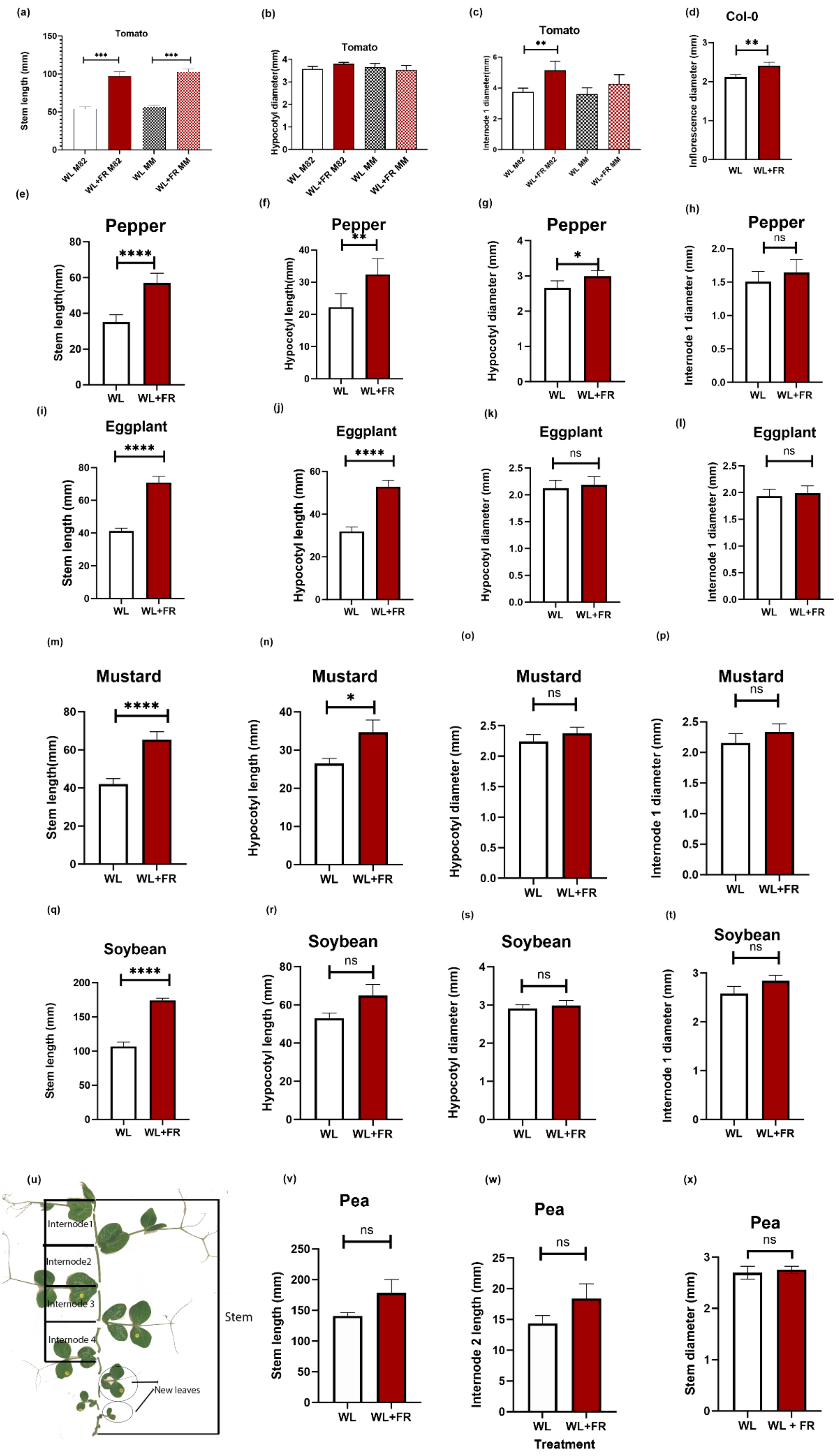

(a)
Tomato
Stem length (mm)
***
***
WL M82
WL+FR M82
WL MM
WL+FR MM
(b)
Tomato
Hypocotyl diameter(mm)
WL M82
WL+FR M82
WL MM
WL+FR MM
(c)
Tomato
Internode 1 diameter(mm)
**
WL M82
WL+FR M82
WL MM
WL+FR MM
(d)
Col-0
Inflorescence diameter (mm)
**
WL
WL+FR
(e)
Pepper
Stem length(mm)
****
WL
WL+FR
(f)
Pepper
Hypocotyl length(mm)
**
WL
WL+FR
(g)
Pepper
Hypocotyl diameter (mm)
*
WL
WL+FR
(h)
Pepper
Internode 1 diameter (mm)
ns
WL
WL+FR
(i)
Eggplant
Stem length (mm)
****
WL
WL+FR
(j)
Eggplant
Hypocotyl length (mm)
****
WL
WL+FR
(k)
Eggplant
Hypocotyl diameter (mm)
ns
WL
WL+FR
(l)
Eggplant
Internode 1 diameter (mm)
ns
WL
WL+FR
(m)
Mustard
Stem length(mm)
****
WL
WL+FR
(n)
Mustard
Hypocotyl length (mm)
*
WL
WL+FR
(o)
Mustard
Hypocotyl diameter (mm)
ns
WL
WL+FR
(p)
Mustard
Internode 1 diameter (mm)
ns
WL
WL+FR
(q)
Soybean
Stem length (mm)
****
WL
WL+FR
(r)
Soybean
Hypocotyl length (mm)
ns
WL
WL+FR
(s)
Soybean
Hypocotyl diameter (mm)
ns
WL
WL+FR
(t)
Soybean
Internode 1 diameter (mm)
ns
WL
WL+FR
(u)
Internode1
Internode2
Internode 3
Internode 4
Stem
New leaves
(v)
Pea
Stem length (mm)
ns
WL
WL+FR
(w)
Pea
Internode 2 length (mm)
ns
WL
WL+FR
Treatment
(x)
Pea
Stem diameter (mm)
ns
WL
WL + FR

**Figure S4: Shoot traits measured in multiple dicot species under WL and WL+FR treatments.** This figure displays (a) stem length, (b) hypocotyl length, (c) first internode diameter of MM and M82 tomato cultivars, (d) Col-0 inflorescence diameter; for bell pepper (e) stem length, (f) hypocotyl length, (g) hypocotyl diameter, (h) first internode diameter; for eggplant (i) stem length, (j) hypocotyl length, (k) hypocotyl diameter, (l) first internode diameter; for soybean (q) stem length, (r) hypocotyl length, (s) hypocotyl diameter, (t) first internode diameter; and for pea (u) structural morphology, (v) stem length, (w) second internode length, (x) stem diameter in WL vs. WL+FR. Significance of differences between treatments is indicated by asterisks: *$p \leqslant 0.05$; **$p \leqslant 0.01$; ***$p \leqslant 0.001$. Error bars denote standard error (SE), with a sample size of n=18 for each measurement.

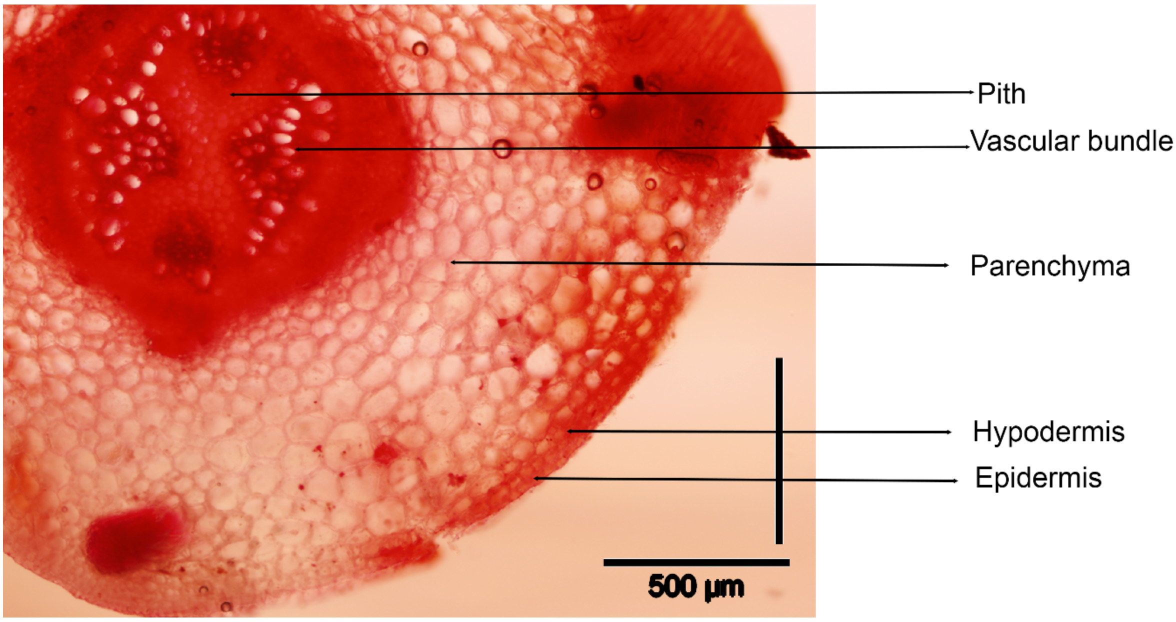


**Figure S5: Cross section of pea (*P. sativum)* internode and cell type identification.**

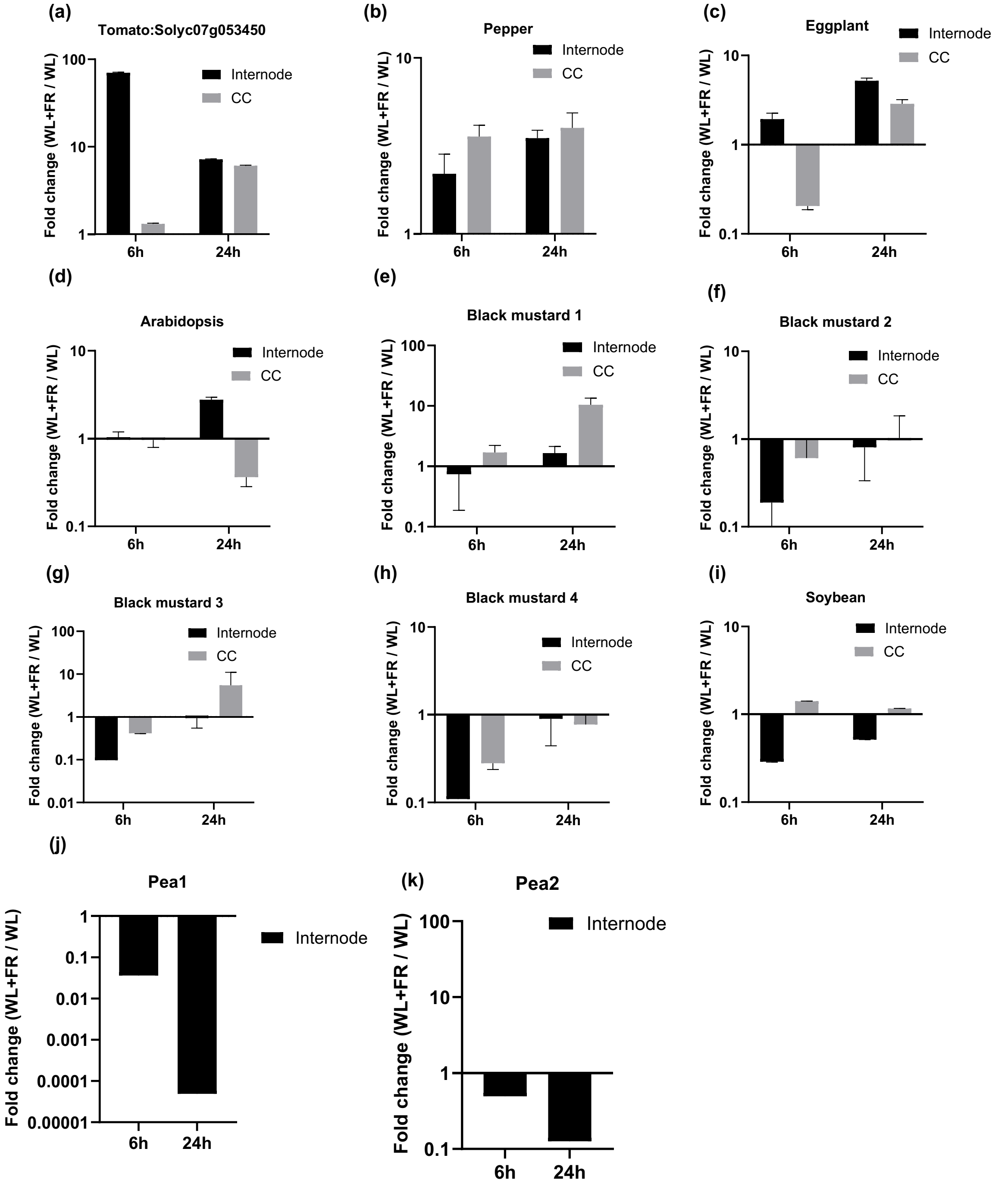


**Figure S6: Fold change of transcript abundance of *Solyc07g053450* homologs in response to FR treatment.** This analysis, conducted using qPCR, focused on the transcript abundance in internode 1 and its central cylinder (CC) at two distinct time points: 6 hours and 24 hours. We compared the gene expression response under WL and WL+FR conditions for the following homologous genes in multiple species: ((a) Tomato: *Solyc07g053450*, (b) Pepper: *CA00g71840:1-921*, (c) Eggplant: *SMEL4.1_07g019740.1.01*, (d) Arabidopsis: *AT2G42380*, (e) Mustard1: *BniB06g001480.2N.1*, (f) Mustard2: *BniB08g027470.2N.1*, (g) Mustard3: *BniB08g072820.2N.1*, (h) Mustard4: *BniB06g055220.2N.1*, (i) Soybean: *Gm06:49537899..49539195*, (j) Pea1: *XM_051027770.1*, (k) Pea2: *XM_051045459.1*. Each sample had 4 biological replicates, comprising 8-12 plants each.

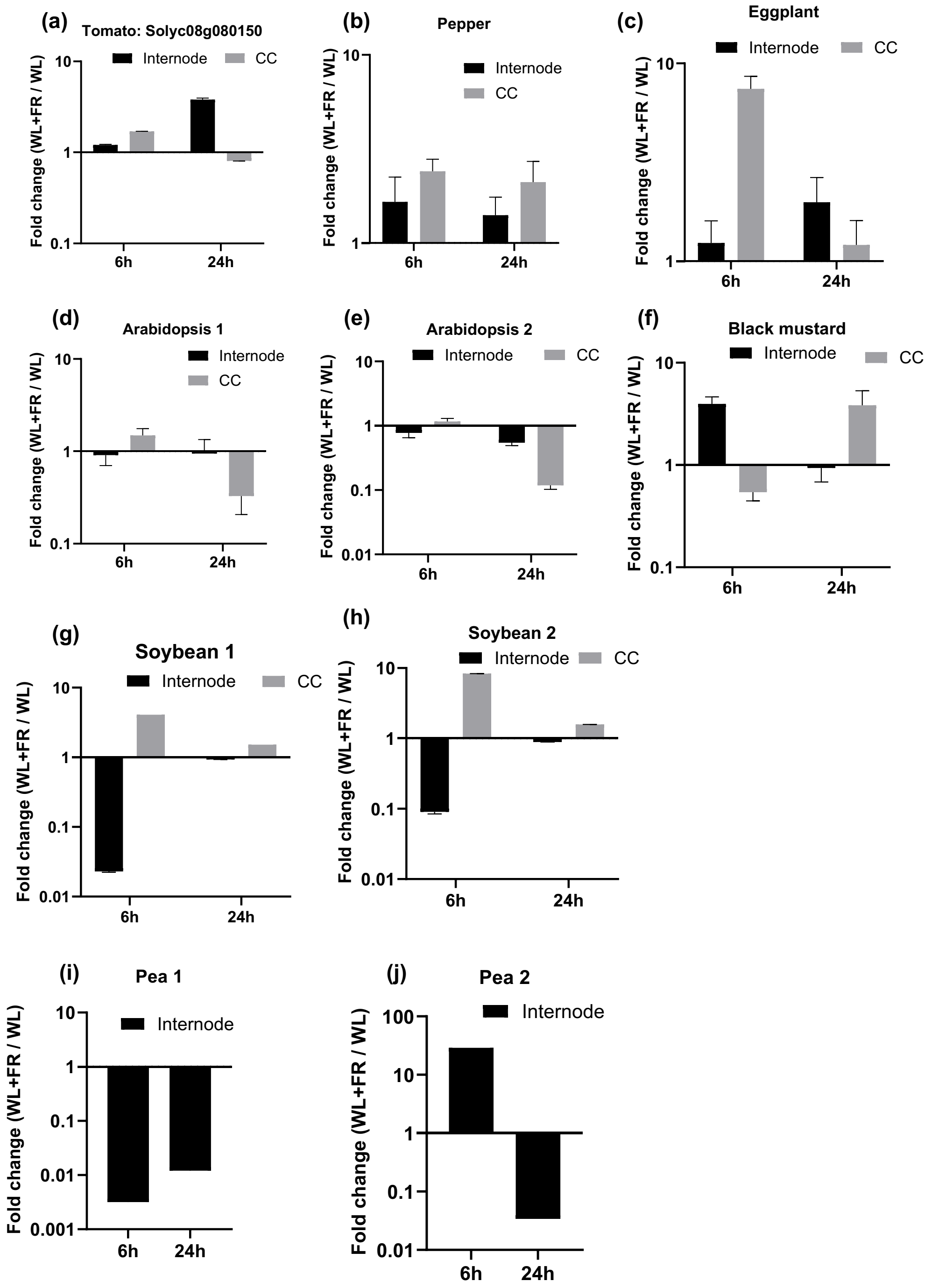


**Figure S7: Fold change of transcript abundance of *Solyc08g080150* homologs in response to FR treatment.** This analysis, conducted using qPCR, focused on the transcript abundance in internode 1 and its central cylinder (CC) at two distinct time points: 6 hours and 24 hours. We compared the gene expression response under WL and WL+FR conditions for the following homologous genes in multiple species: (a) Tomato: *Solyc08g080150*, (b) Pepper: *TCP19*, (c) Eggplant: *chr8:85852001-85853900*, (d) Arabidopsis1: *AT2G45680*, (e) Arabidopsis2: *AT5G51910*, (f) Mustard: *BniB08g029800.2N.1*, (g) Soybean1: *Glyma.07G080300.1*, (h) Soybean2: *Glyma07g08710.2*, (i) Pea1: *XM_051030490.1*, (j) Pea2: *XM_051030491.1*. Each sample had 4 biological replicates, comprising 8-12 plants each.

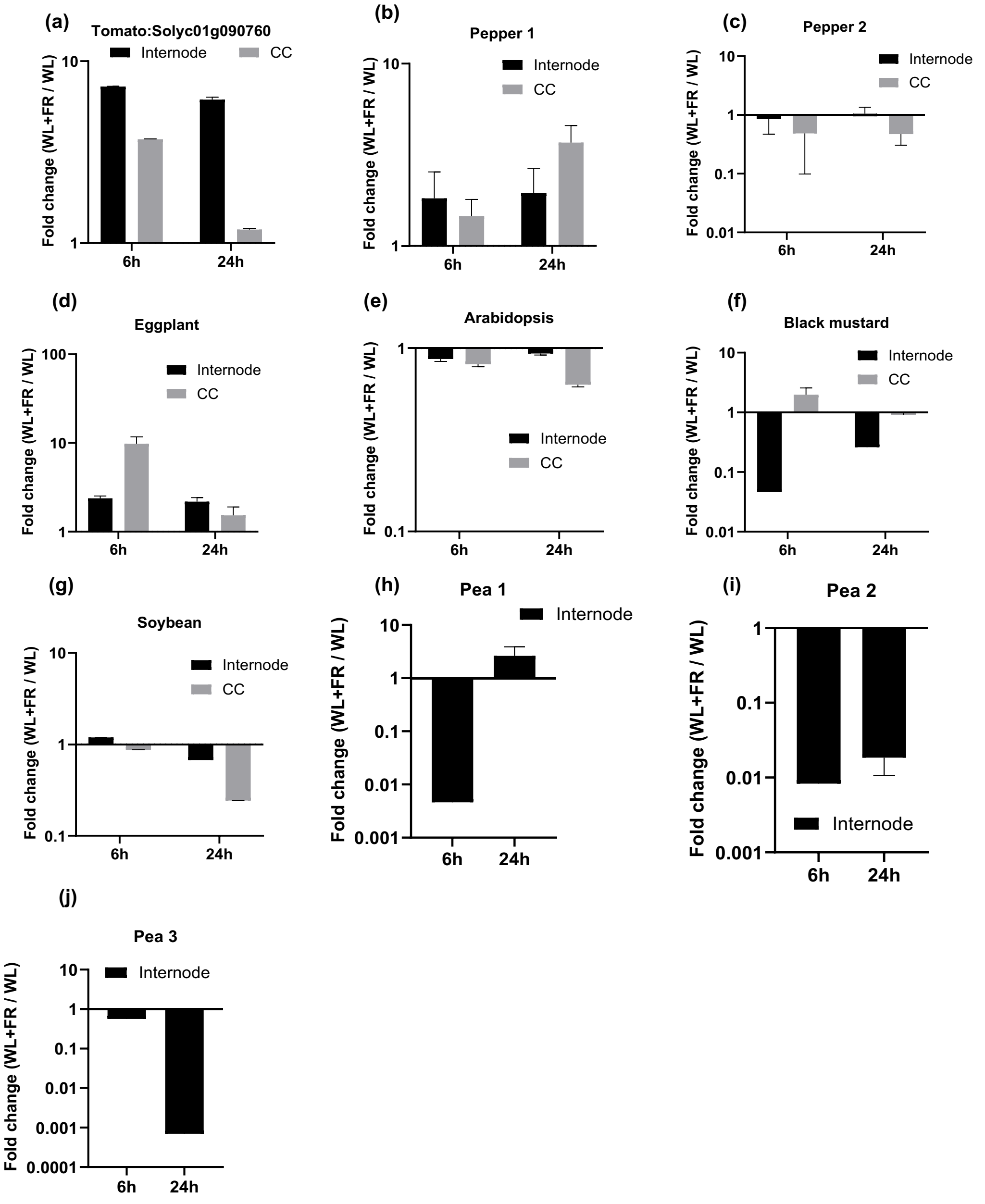


**Figure S8: Fold change of transcript abundance of *Solyc01g090760* homologs in response to FR treatment.** This analysis, conducted using qPCR, focused on the transcript abundance in internode 1 and its central cylinder (CC) at two distinct time points: 6 hours and 24 hours. We compared the gene expression response under WL and WL+FR conditions for the following homologous genes in multiple species: (a) Tomato: *Solyc01g090760*, (b) Pepper1: *chr3:267174801-267176100* (*CA00g71840:1-921*), (c) Pepper2*: chr8:35902201-35904000*, (d) Eggplant: *chr8:35902201-35904000*, (e) Arabidopsis: *AT2G45050*, (e) Mustard: *BniB07g039570.2N.1*, (f) Soybean: *GlysoPI483463.06G078800.1*, (g) Pea1: *XM_051027377*, (h) Pea2: *XM_051030552.1*, (i) Pea3: *XM_051038772.1*. Each sample had 4 biological replicates, comprising 8-12 plants each.
